# Acyl-CoA Binding Protein in White and Brown Adipose Tissue is Dispensable for Systemic Energy Metabolism

**DOI:** 10.1101/2025.04.10.642978

**Authors:** M. F. Nørremark, R. Petersen, P. M. M. Ruppert, E. S. Jul, T. K. Doktor, R. Nielsen, J. F. Kappel, S. Larsen, J. W. Kornfeld, S. Mandrup, B. S. Andresen, J. Havelund, D. Neess, N. J. Færgeman

## Abstract

Acyl-CoA binding protein (ACBP) plays a vital role in lipid metabolism by mediating the intracellular flux and utilization of long-chain acyl-CoAs. In this study we generated brown- and white adipose tissue specific knockout mice (Adipoq-*Acbp^-/-^*) and brown adipose tissue specific knockout mice (Ucp1-*Acbp^-/-^*) to investigate the role of ACBP in adipose tissue function. Here we demonstrate that loss of ACBP does not affect body weight, fat and lean mass, food intake and systemic energy expenditure, even under cold stress. Transcriptomic data show only minor changes in gene expression, whereas lipidomic profiling reveals a subtle increase in acyl-carnitines levels in brown adipose tissue. However, lipolytic activity in white adipose tissue as well as plasma glycerol, non-esterified fatty acid and triacylglycerol levels remained unaffected. In addition, no changes in mitochondrial respiration in BAT were observed. Taken together, our findings suggest that ACBP is dispensable for adipose tissue function and systemic energy metabolism, including thermoregulation.

## Introduction

Adipose tissues exhibit a high degree of plasticity, making them essential players in the regulation of systemic energy homeostasis. A remarkable feature of adipose tissue is the ability to undergo rapid and dynamic remodeling to adapt to variations in nutrient availability and environmental conditions(1). Traditionally, adipose tissue has been categorized into white adipose tissue (WAT) and brown adipose tissue (BAT), but a third type, beige (or brite) adipose tissue, is now recognized(2). The distinct brown color of BAT is attributed to the high density of mitochondria in the multilocular brown adipocytes. BAT is a thermogenic tissue that contributes to non-shivering thermogenesis (NST) through heat generation fueled by lipid oxidation(1, 3). WAT serves as a caloric reservoir, critical for energy storage and insulation and is primarily comprised of unilocular white adipocytes(4). Beige adipocytes, found within WAT depots, can acquire thermogenic properties in response to stimuli such as cold or β-adrenergic activation(5). The energy is stored in intracellular lipid droplets as triacylglycerols (TAGs). TAG synthesis primarily occurs in liver and adipose tissues(6), where fatty acyl-chains are esterified with glycerol 3-phosphate in a multistep enzymatic process(7). Most of the fatty acids used for TAG synthesis are provided by circulating plasma lipids, such as non-esterified fatty acids (NEFA) and lipoprotein-derived TAGs, or through the complex process of *de novo* lipogenesis (DNL), which is the synthesis of new fatty acids from non-lipid substrates, mainly carbohydrates. When energy demand is high or nutrients are scarce, TAGs undergo lipolysis generating glycerol and free fatty acids, which are transported via the circulation to utilizing tissues incl. muscle and liver(8). However, fatty acids cannot be metabolized without prior activation to their CoA derivatives, catalyzed by the family of acyl-CoA synthetases(9). Once activated, acyl-CoAs can be oxidized to acetyl-CoA to release energy or be incorporated into more complex lipids such as glycerolipids, phospholipids and sphingolipids(10). The intracellular concentration and selective partitioning of acyl-CoAs are tightly controlled by transport proteins, such as acyl-CoA binding protein (ACBP)(10). ACBP is a 10 kDa cytosolic protein, which specifically binds medium-, long- and very-long-chain fatty acyl-CoA esters with high affinity(11, 12). ACBP was originally identified as diazepam binding inhibitor (DBI) due to its ability to modulate benzodiazepine binding to the GABA_A_ receptor(13), however, ACBP has also been recognized for its crucial roles in intracellular lipid metabolism. *In vitro*, ACBP prevents the inhibitory effect that long-chain fatty acyl-CoA (LCACoAs) exert on enzymes such as the mitochondrial acyl-CoA synthetase, acetyl-CoA carboxylase as well as the mitochondrial nucleotide translocase(14). Furthermore, by donating LCACoAs to utilizing enzymes, ACBP stimulates synthesis of glycerolipids, glycerophospholipids, very-long-chain fatty acids, and ceramides. ACBP also promotes mitochondrial β-oxidation(15).

Recent *in vivo* studies have shown that ACBP is required for proper development and maintenance of the epidermal barrier(12, 16, 17). Genetic ablation of *Acbp* in mice results in increased trans-epidermal water loss and heat dissipation, as well as development of alopecia and depigmentation of the fur. These phenotypes are recapitulated in mice lacking ACBP due to targeted disruption in keratinocytes(12). Both systemic loss of *Acbp* and specific loss in keratinocytes results in increased energy expenditure, increased food intake and increased browning of the inguinal white adipose tissue (iWAT), which all are reversed by housing the mice at 30°C or by blocking β-adrenergic receptor signalling(12). Consistent with increased energy expenditure, both full-body and keratinocyte-specific *Acbp*^-/-^ mice are completely resistant to diet-induced obesity and the diabetogenic effects of a high-fat diet(12). Noticeably, *Acbp*^-/-^ mice display a higher respiratory quotient than wild type and keratinocyte-specific *Acbp*^-/-^ mice, arguing that ACBP is likely to facilitate oxidation of fatty acids in other cell types than keratinocytes, and thereby contribute to systemic utilization of fatty acids as energy substrates(12). These findings prompted us to examine the function of ACBP in adipose tissues. Here we demonstrate that mice with tissue-specific loss of ACBP in white and/or brown adipose tissue exhibit no changes in body weight and temperature, food intake, body mass distribution and weight of selected tissues during both standard and cold housing conditions. In addition, we show that lack of ACBP in adipose tissues only induce limited changes in the transcriptome, metabolic phenotype and mitochondrial respiration capacity. Notably, we report elevated levels of acyl-carnitines in BAT from Adipoq-*Acbp^-/-^* mice housed at room temperature. However, we find no evidence of alterations in lipolytic activity or substrate flux from the white adipose tissue. Collectively, these data show that ACBP is dispensable for the adipose tissue function in mice.

## Results

### Tissue-specific ablation of *Acbp* in adipose tissues

To investigate the intracellular function of ACBP in adipose tissues, we generated two mouse strains with conditional targeting of the *Acbp* gene in adipocytes. These models were generated by mating mice carrying a floxed *Acbp* allele with mice expressing the Cre recombinase under control of either the Adiponectin promoter (Adipoq-Cre) or Uncoupling protein 1 promoter (Ucp1-Cre). Mating with Adipoq-Cre mice generates a brown- and white adipocyte-specific knockout of ACBP (Adipoq-*Acbp^-/-^*), whereas mating with Ucp1-Cre mice results in a brown adipocyte-specific knockout of ACBP (Ucp1-*Acbp^-/-^*). To assess the extent of *Acbp* knockout in different adipose depots, we initially analyzed the mRNA levels of *Acbp* using quantitative rt-PCR. The *Acbp* mRNA expression was not affected in liver and muscle tissue from Adipoq-*Acbp^-/-^* mice but was significantly reduced compared with controls in several adipose tissues, including gonadal WAT (gWAT), inguinal WAT (iWAT) and interscapular brown adipose tissue (BAT) (Figure 1A). As expected, we found a significant reduction in *Acbp* mRNA levels only in BAT of Ucp1-*Acbp^-/-^* mice, whereas *Acbp* expression in liver, muscle, gWAT and iWAT was comparable with that of control mice (Figure 1B). Accordingly, the ACBP protein was not detectable in iWAT and BAT of Adipoq-*Acbp^-/-^* mice (Figure 1C) and similarly not detectable in BAT of Ucp1-*Acbp^-/-^* mice (Figure 1D). ACBP protein levels in liver were unaffected for both strains (Figure 1C-D). Additionally, immunohistochemical detection of ACBP demonstrated that adipocytes in iWAT and BAT from Adipoq-*Acbp^-/-^*mice were depleted of ACBP (Figure 1E), whereas only adipocytes in BAT from Ucp1-*Acbp^-/-^* mice lacked ACBP expression (Figure 1F). Collectively, these data show efficient and specific knockout of ACBP in iWAT and BAT of Adipoq-*Acbp^-/-^* mice as well as BAT of Ucp1-*Acbp^-/-^* mice.

**Figure 1.**
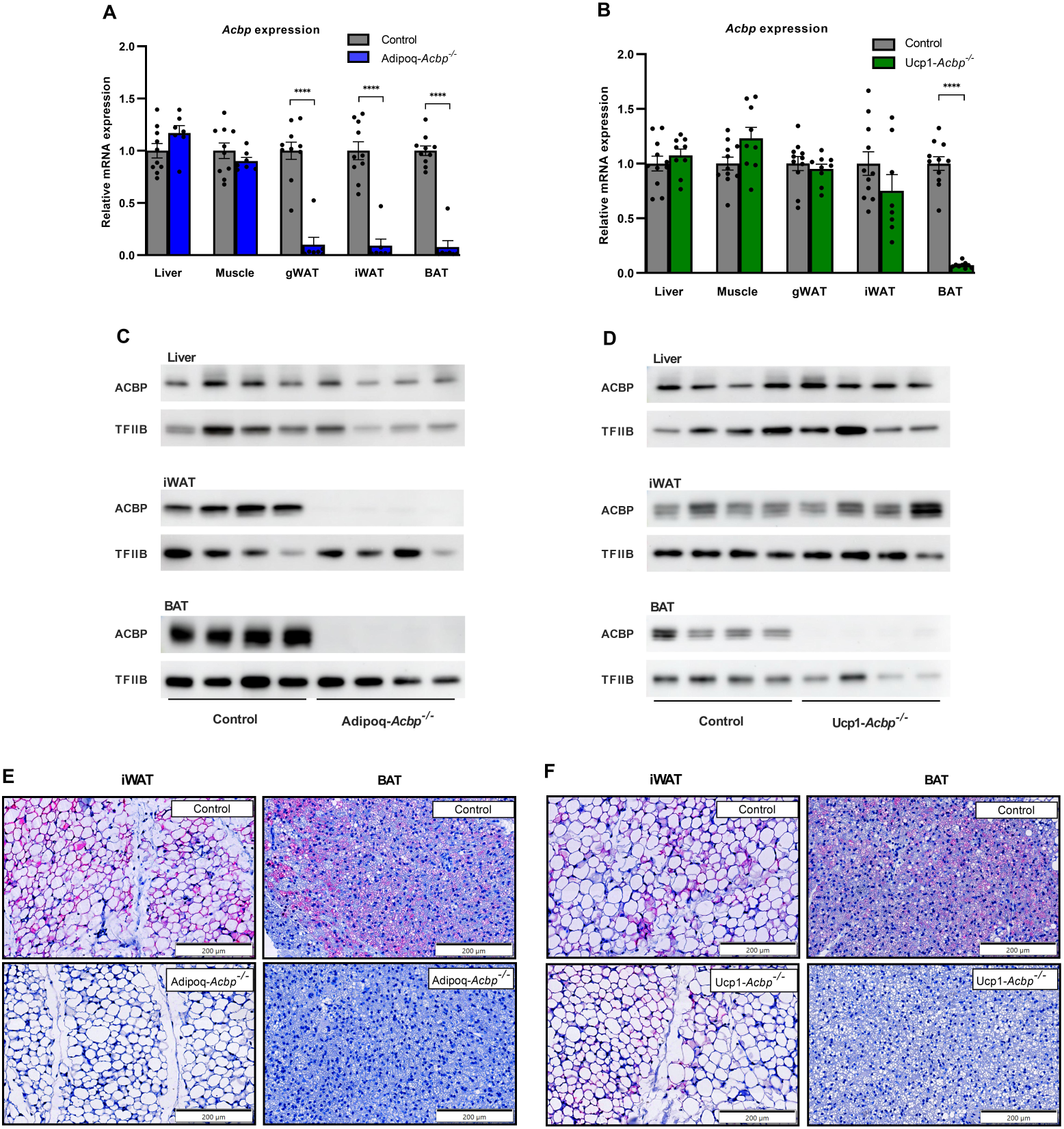
Tissue specific ablation of *Acbp* in adipose tissues. (A) mRNA levels of *Acbp* in liver, muscle (quadriceps), gWAT, iWAT and BAT harvested from control and Adipoq-*Acbp^-/-^* mice housed at 22°C. (n=7-10 per group, Two-way ANOVA). Data are presented as mean of individuals in each group ± SEM. ****p<0.0001. (B) mRNA levels of *Acbp* in liver, muscle (quadriceps), gWAT, iWAT and BAT harvested from control and Ucp1-*Acbp^-/-^* mice housed at 22°C. (n=9-11 per group, Two-way ANOVA). Data are presented as mean of individuals in each group ± SEM. ****p<0.0001. (C) ACBP expression determined by western blotting of liver, iWAT and BAT harvested from control and Adipoq-*Acbp^-/-^* mice housed at 22°C. For each tissue, protein extracts from four individual mice were analyzed by western blotting and probed for ACBP and TFIIB. (D) ACBP expression determined by western blotting of liver, iWAT and BAT harvested from control and Ucp1-*Acbp^-/-^* mice housed at 22°C. For each tissue, protein extracts from four individual mice were analyzed by western blotting and probed for ACBP and TFIIB. (E) Representative ACBP immunostaining images of iWAT and BAT harvested from control (upper panel) and Adipoq-*Acbp^-/-^* (lower panel) mice housed at 22°C. Scale bars are 50 µm and each image represents 10 individuals per group. (F) Representative ACBP immunostaining images of iWAT and BAT harvested from control (upper panel) and Ucp1-*Acbp^-/-^*(lower panel) mice housed at 22°C. Scale bars are 50 µm and each image represents 10 individuals per group. Mice were 10-14 weeks old. All mice were sacrificed by cervical dislocation.

### Loss of ACBP in adipose tissues does not affect body- and tissue weights and food intake

To explore whether loss of ACBP in adipose tissues impacts the size and body composition of the mice, we examined their body weight, food intake, body mass distribution and weight of selected tissues. Notably, we found no significant changes in body weight and food intake for either Adipoq-*Acbp^-/-^* mice or Ucp1-*Acbp^-/-^* mice compared with the controls (Figure 2A and 2B, respectively). TD-NMR was applied to examine the body composition of Adipoq-*Acbp^-/-^*and Ucp1-*Acbp^-/-^* mice. However, we found no differences in lean mass and body fluid levels between mice of the different genotypes (Figure 2C and 2D). To examine whether ACBP depleted adipose tissues developed differently, we determined the weight of iWAT and BAT tissues, but the data revealed no differences in tissue weight between Adipoq-*Acbp^-/-^* or Ucp1-*Acbp^-/-^*mice and their respective controls when normalized to body weight (Figure 2E and 2F). This prompted us to examine if loss of ACBP would alter the adipogenic potential of stroma vascular fractions (SVF) from WAT and BAT. Thus, we examined the expression of key adipogenic markers of primary white and brown preadipocytes from wildtype or *Acbp^-/-^* mice during differentiation. In line with the results above, loss of ACBP did not affect expression of WAT and BAT markers arguing that ACBP is not required for differentiation of primary preadipocytes to mature adipocytes (Figure S1).

**Figure 2.**
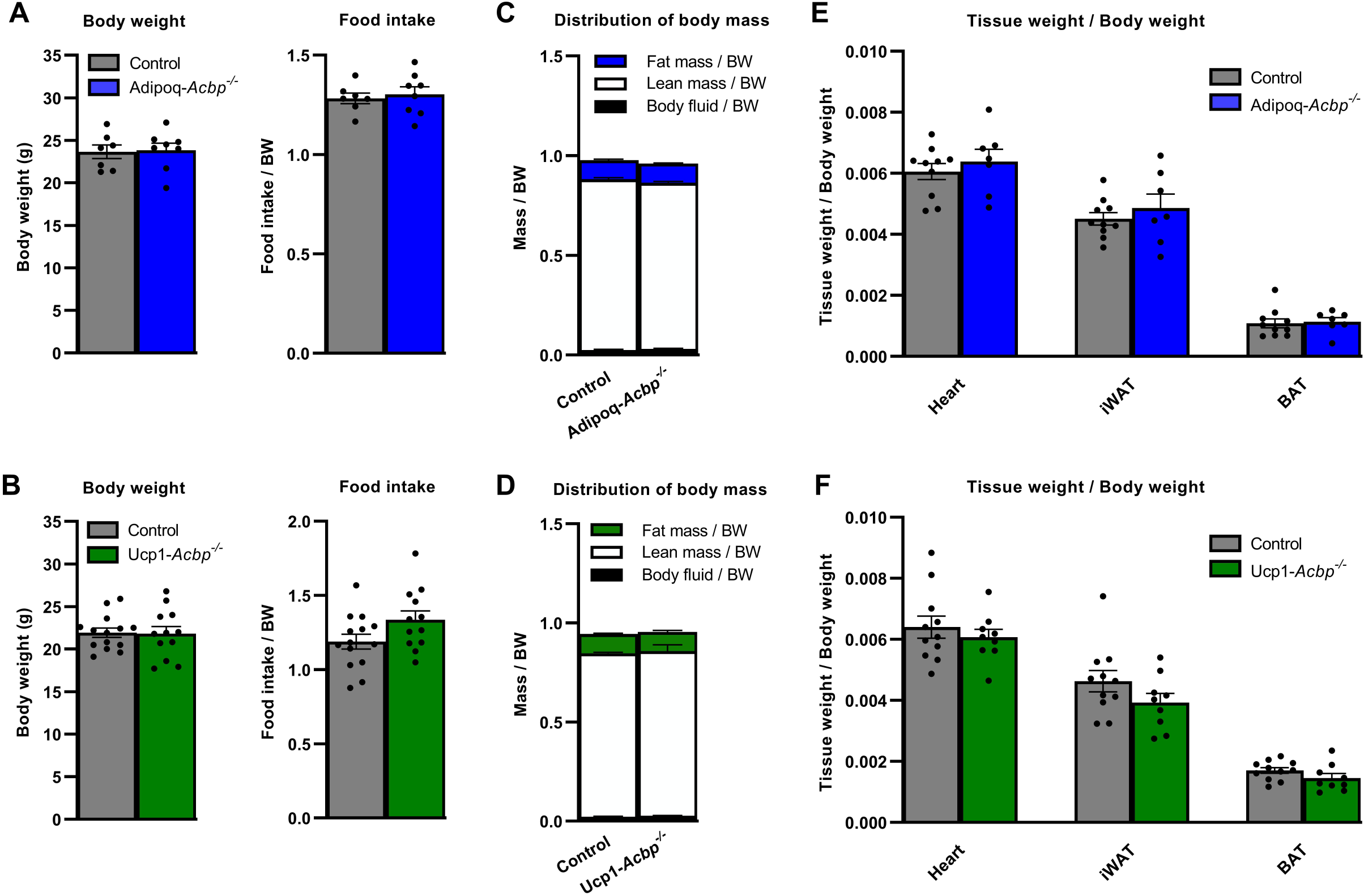
Loss of ACBP in adipose tissues does not affect body- and tissue weights and food intake at room temperature. (A) Control and Adipoq-*Acbp^-/-^*mice were housed individually for 7 days at 22°C. The food intake was monitored and after 7 days the body weight of all mice was determined. (n= 7-8 per group, unpaired t-test). Data for body weight are presented as mean of individuals in each group ± SEM. Data for food intake are presented relative to body weight (BW) as mean of individuals in each group ± SEM. (B) Control and Ucp1-*Acbp^-/-^* mice were housed individually for 7 days at 22°C. The food intake was monitored and after 7 days the body weight of all mice was determined. (n=12-14 per group, unpaired t-test). Data for body weight are presented as mean of individuals in each group ± SEM. Data for food intake are presented relative to body weight (BW) as mean of individuals in each group ± SEM. (C) Control and Adipoq-*Acbp^-/-^* mice were housed individually for 7 days at 22°C. After 7 days, the body composition was analyzed using NMR scanning (MiniSpec LF50). Data for fat mass, lean mass and body fluid relative to body weight (BW) are presented as mean of individuals in each group ± SEM. (n=12-16 per group, Two-way ANOVA). (D) Control and Ucp1-*Acbp^-/-^* mice were housed individually for 7 days at 22°C. After 7 days, the body composition was analyzed using NMR scanning (MiniSpec LF50). Data for fat mass, lean mass and body fluid relative to body weight (BW) are presented as mean of individuals in each group ± SEM. (n=12-14 per group, Two-way ANOVA). (E) Control and Adipoq-*Acbp^-/-^* mice were housed individually for 7 days at 22°C. After 7 days, the mice were sacrificed and the heart, iWAT and BAT were weighed during dissection. Data for tissue weight relative to body weight are presented as mean of individuals in each group ± SEM. (n=7-10 per group, Two-way ANOVA). (F) Control and Ucp1-*Acbp^-/-^* mice were housed individually for 7 days at 22°C. After 7 days, the mice were sacrificed and the heart, iWAT and BAT were weighed during dissection. Data for tissue weight relative to body weight are presented as mean of individuals in each group ± SEM. (n=9-11 per group, Two-way ANOVA). Mice were 10-14 weeks old. In Figure A-D, mice were anesthetized for 5 minutes and subsequently sacrificed by heart puncture to draw blood. In Figure E and F mice were sacrificed by cervical dislocation.

Cold exposure is a well-established method to stimulate thermogenic responses and mobilize energy stores(18). To determine whether ACBP deficiency in adipose tissues influence the adaptive response to cold stress, we subjected Adipoq-*Acbp^-/-^* and Ucp1-*Acbp^-/-^* to 4°C for 7 days to assess whether cold exposure could lead to phenotypic differences in body composition or tissue distribution. However, cold exposure did not induce alterations in body weight or food intake in Adipoq-*Acbp^-/-^*and Ucp1-*Acbp^-/-^* mice compared with their control littermates (Figure 3A and 3B). No effects were observed on body mass distribution (Figure 3C and 3D) and size of adipose tissues for both mouse models (Figure 3E and 3F). Notably, under cold conditions, Adipoq-*Acbp^-/-^* and Ucp1-*Acbp^-/-^* mice were able to maintain a normal core body temperature similar to what was observed for the controls, even in the absence of food for six hours (Figure 3G and 3H). In summary, these data demonstrate, that mice can defend their body temperature and maintain normal tissue distribution independent of the presence of ACBP in adipose tissues.

**Figure 3.**
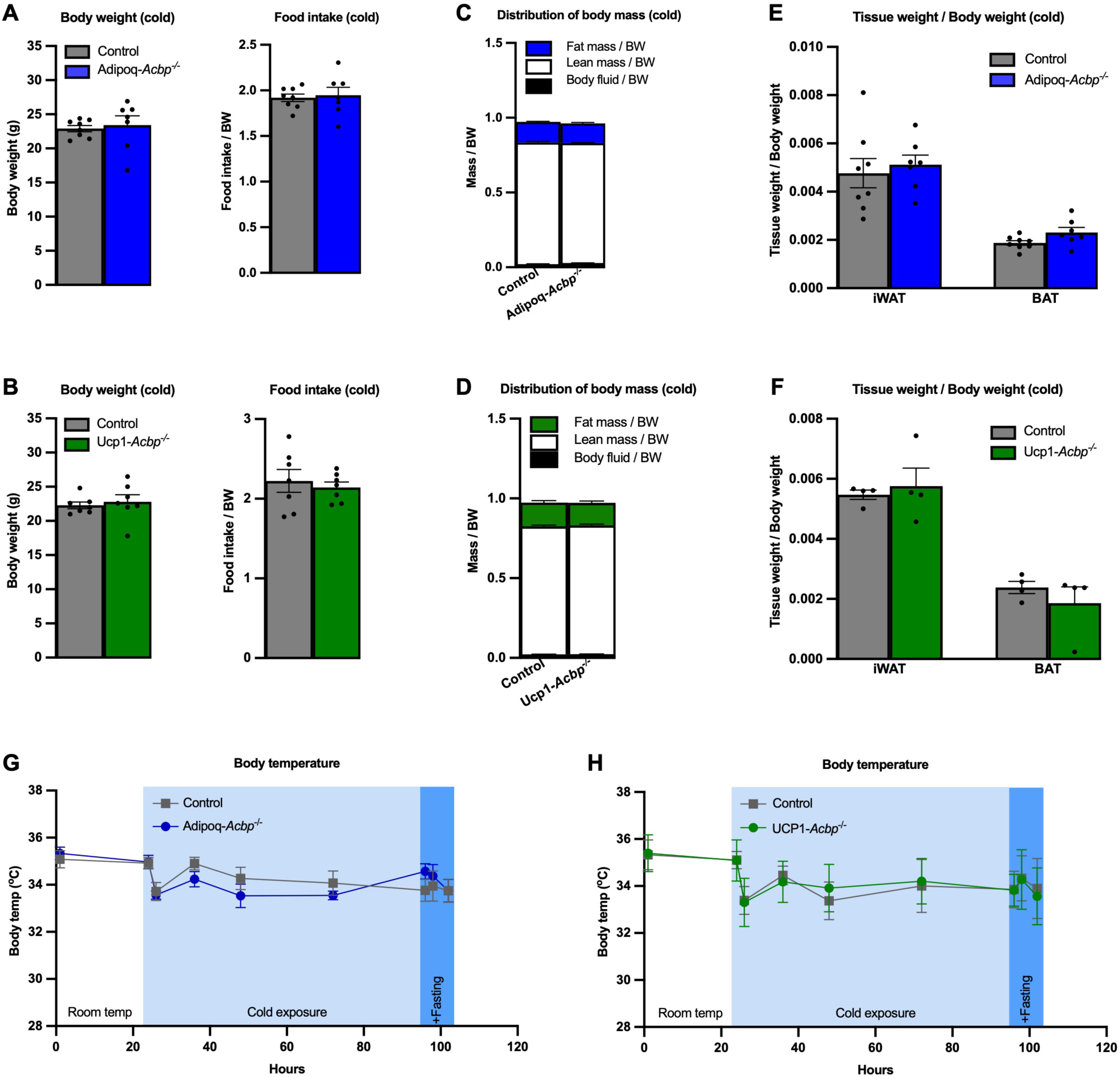
Loss of ACBP in adipose tissues does not affect body- and tissue weights and food intake during cold exposure. (A) Control and Adipoq-*Acbp^-/-^* mice were housed individually for 7 days at 4°C. The food intake was monitored and after 7 days the body weight of all mice was determined (n=7-8 per group, unpaired t-test). Data for body weight are presented as mean of individuals in each group ± SEM. Data for food intake are presented relative to body weight (BW) as mean of individuals in each group ± SEM. (B) Control and Ucp1-*Acbp^-/-^* mice were housed individually for 7 days at 4°C. The food intake was monitored and after 7 days the body weight of all mice was determined (n=7 per group, unpaired t-test). Data for body weight are presented as mean of individuals in each group ± SEM. Data for food intake are presented relative to body weight (BW) as mean of individuals in each group ± SEM. (C) Control and Adipoq-*Acbp^-/-^* mice were housed individually for 7 days at 4°C. After 7 days, the body composition was analyzed using NMR scanning (MiniSpec LF50). Data for fat mass, lean mass and body fluid relative to body weight (BW) are presented as mean of individuals in each group ± SEM. (n=5-10 per group, Two-way ANOVA). (D) Control and Ucp1-*Acbp^-/-^* mice were housed individually for 7 days at 4°C. After 7 days, the body composition was analyzed using NMR scanning (MiniSpec LF50). Data for fat mass, lean mass and body fluid relative to body weight (BW) are presented as mean of individuals in each group ± SEM. (n=7 per group, Two-way ANOVA). (E) Control and Adipoq-*Acbp^-/-^* mice were housed individually for 7 days at 4°C. The heart, iWAT and BAT were weighed during dissection. Data for tissue weight relative to body weight are presented as mean of individuals in each group ± SEM. (n=7-8 per group, Two-way ANOVA). (F) Control and Ucp1-*Acbp^-/-^* mice were housed individually for 7 days at 4°C. The heart, iWAT and BAT were weighed during dissection. Data for tissue weight relative to body weight are presented as mean of individuals in each group ± SEM. (n=4 per group, Two-way ANOVA). (G) The body temperature of control and Adipoq-*Acbp^-/-^*mice was monitored by implantation of subcutaneous transponders and the mice were subsequently acclimatized for 7 days. The temperature was measured at room temperature (22°C) and during acute and chronic cold exposure (4°C). Data are presented as mean of individuals in each group ± SEM. (n=6 per group, Two-way ANOVA). (H) The body temperature of control and Ucp1-*Acbp^-/-^* mice was monitored by implantation of subcutaneous transponders and the mice were subsequently acclimatized for 7 days. The temperature was measured at room temperature (22°C) and during acute and chronic cold exposure (4°C). Data are presented as mean of individuals in each group ± SEM. (n=6-7 per group, Two-way ANOVA). Mice were 7-12 weeks old. In Figure A-F, mice were anesthetized for 5 minutes and subsequently sacrificed by heart puncture to draw blood. In Figure G and H mice were sacrificed by cervical dislocation.

### Loss of ACBP in adipose tissues leads to minor transcriptional changes

To identify pathways affected by functional loss of ACBP in adipose tissue, we examined the transcriptional changes in iWAT, and BAT induced by ablation of the *Acbp* gene. We therefore performed RNA sequencing (RNA-seq) on iWAT and BAT from Adipoq-*Acbp^-/-^* mice and BAT from Ucp1-*Acbp^-/-^* mice housed at room temperature (RT). Differential gene expression analysis confirmed that *Acbp*/*Dbi* as expected, was significantly downregulated in iWAT and BAT from Adipoq-*Acbp^-/-^*mice and BAT from Ucp1-*Acbp^-/-^* mice compared with their controls (Figure 4A-C), further substantiating the data presented in Figure 1. However, RNAseq analyses demonstrated that loss of ACBP in iWAT and BAT in Adipoq-*Acbp^-/-^*mice and BAT in Ucp1-*Acbp^-/-^* mice only induce subtle changes in their gene expression profile compared with their respective controls (Figure 4 and Table S1). These findings show that *Acbp* depletion in adipose tissues only has minor effects on the transcriptome of iWAT and BAT.

**Figure 4.**
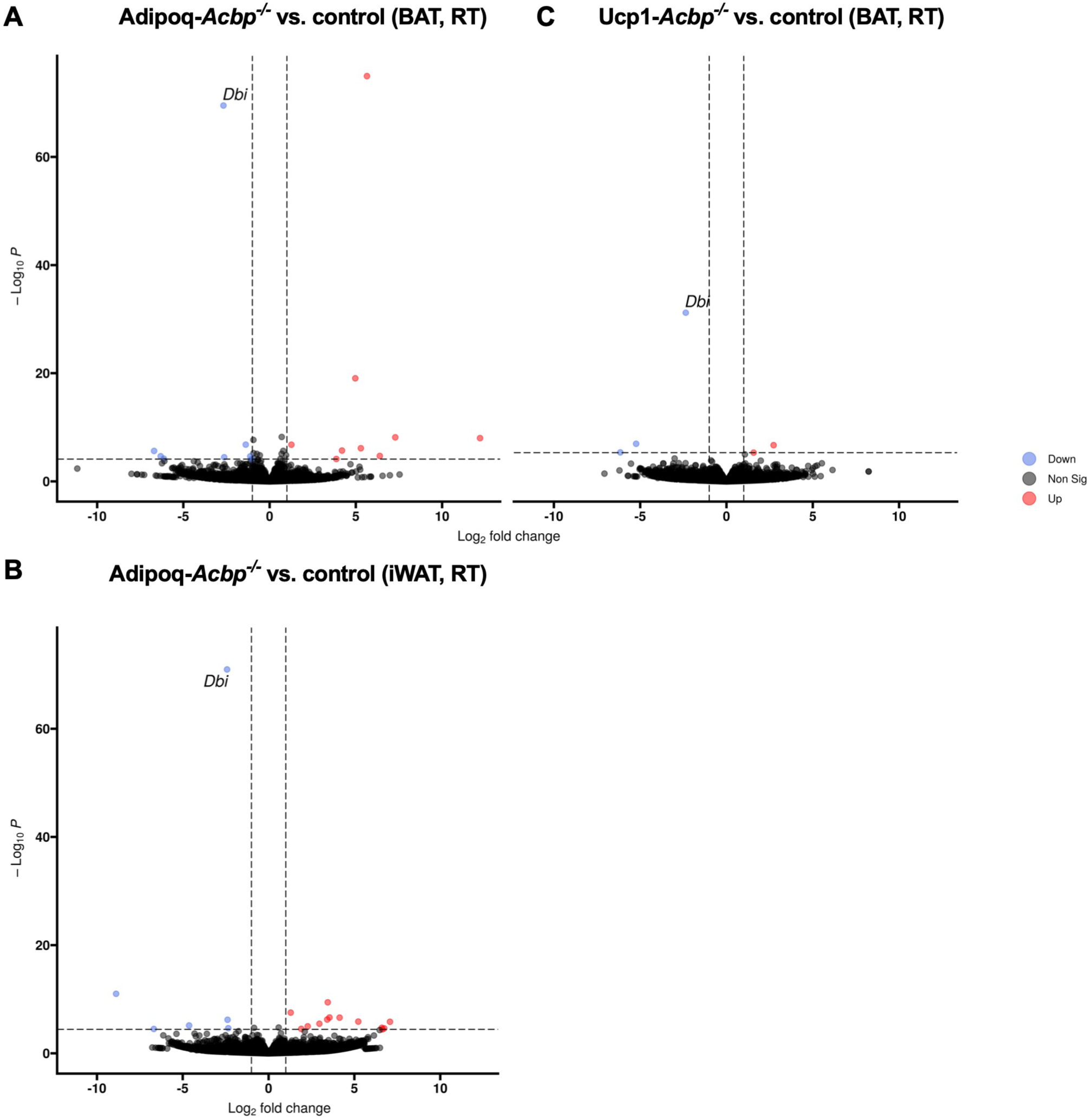
Loss of ACBP in adipose tissues leads to minor transcriptional changes. A) Volcano plot of differentially expressed genes in BAT harvested from Adipoq-*Acbp^-/-^* mice housed individually at room temperature (22°C) for 7 days compared to BAT harvested from control mice housed individually at room temperature (22°C) for 7 days. Red dots indicate significantly upregulated genes. Blue dots indicate significantly downregulated genes. Genes with a p-adjusted value <0.05 and a Log2 fold change >1 were considered as differentially expressed genes (n=3 per group). B) Volcano plot of differentially expressed genes in iWAT harvested from Adipoq-*Acbp^-/-^* mice housed individually at room temperature (22°C) for 7 days compared to iWAT harvested from control mice housed individually at room temperature (22°C) for 7 days. Red dots indicate significantly upregulated genes. Blue dots indicate significantly downregulated genes. Genes with a p-adjusted value <0.05 and a Log2 fold change >1 were considered as differentially expressed genes (n=3 per group). C) Volcano plot of differentially expressed genes in BAT harvested from Ucp1-*Acbp^-/-^*mice housed individually at room temperature (22°C) for 7 days compared to BAT harvested from control mice housed individually at room temperature (22°C) for 7 days. Red dots indicate significantly upregulated genes. Blue dots indicate significantly downregulated genes. Genes with an adjusted p value <0.05 and a Log2 fold change >1 were considered as differentially expressed genes (n=4 per group). Mice were 9-11 weeks old. All mice were sacrificed by cervical dislocation.

### ACBP in adipose tissue is dispensable for systemic energy metabolism

We previously reported that full-body *Acbp^-/-^* and keratinocyte-specific *Acbp^-/-^* mice exhibit increased energy expenditure, increased food intake, increased browning of iWAT and are resistant to diet-induced obesity(12) suggesting that ACBP is a key player in systemic energy homeostasis. We also observed that full-body *Acbp*^-/-^ mice display a higher respiratory quotient than wild type and keratinocyte-specific *Acbp*^-/-^ mice, arguing that ACBP facilitates oxidation of fatty acids in other cell types than keratinocytes. We therefore reasoned that removal of ACBP from the adipose tissue could impair oxidation of fatty acids and hence affect systemic energy homeostasis. Thus, we used indirect calorimetry to assess energy expenditure (EE) and respiratory exchange ratio (RER) of Adipoq-*Acbp^-/-^* and Ucp1-*Acbp^-/-^* mice when housed first at RT (22°C) followed by cold exposure (4°C) and lastly at TN (30°C) for 3 days at each temperature. We found no significant changes in EE, RER as well as food and water intake during the light and dark phase when Adipoq-*Acbp^-/-^* mice (Figure 5A-L) or Ucp1-*Acbp^-/-^* mice (Figure 6A-L) were housed at room temperature, 4°C, or thermoneutrality when compared with their respective control littermates, although there was a slight tendency towards increased energy expenditure in Ucp1-*Acbp^-/-^* mice. Including bodyweight as covariate confirms that loss of ACBP does not affect systemic energy metabolism independent of temperature and genotype (Figure S2). Thus, these data indicate that removal of ACBP from the adipose tissue only have minor effects on systemic energy homeostasis of neither Adipoq-*Acbp^-/-^* mice nor Ucp1-*Acbp^-/-^*mice.

**Figure 5.**
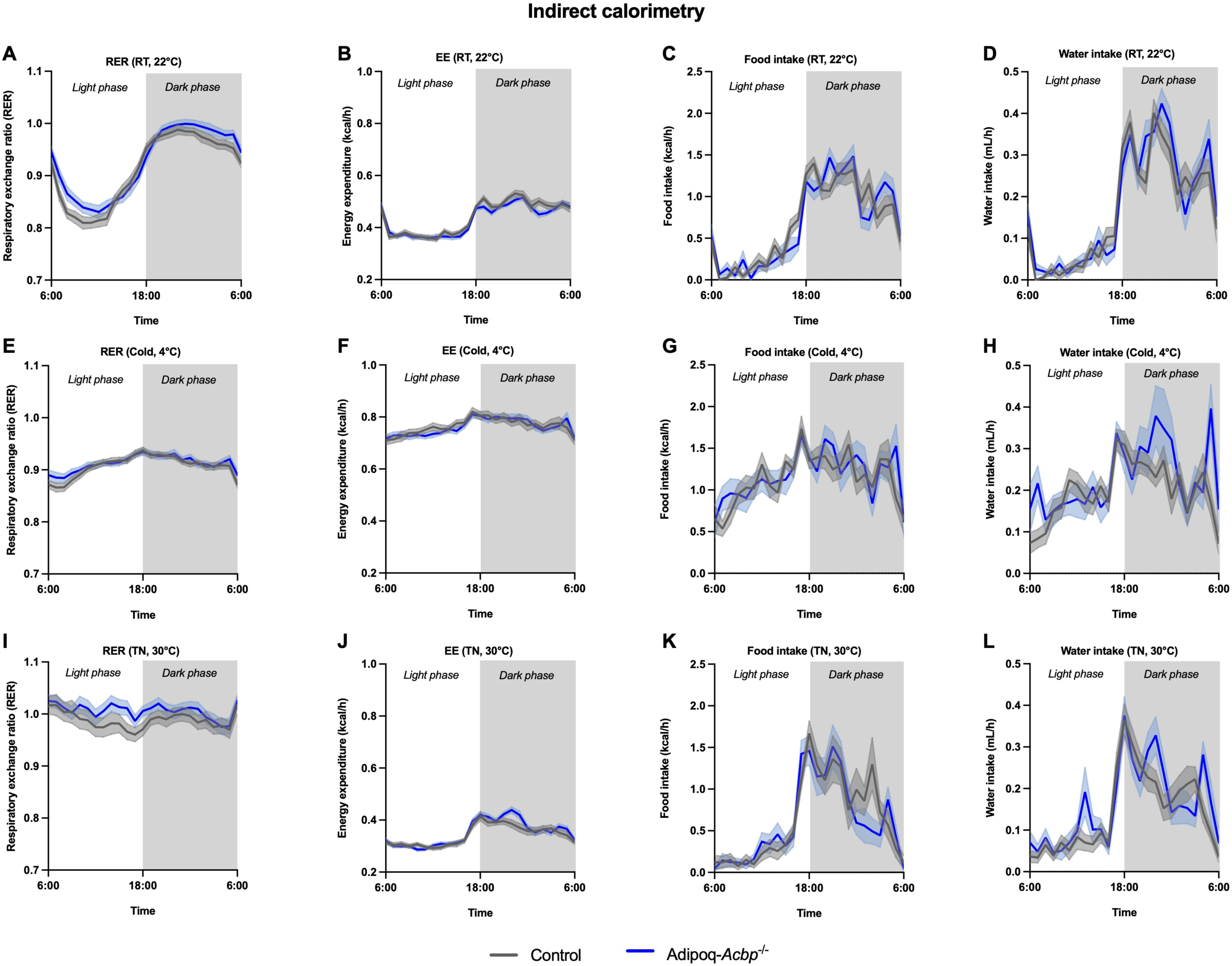
ACBP in adipose tissue is dispensable for systemic energy metabolism in Adipoq-*Acbp^-/-^* mice. A) The respiratory exchange ratio (RER) was examined for Adipoq-*Acbp^-/-^*and control mice housed individually at room temperature (22°C) for 72 hours in metabolic cages. Data are presented as mean of individuals in each group ± SEM (n=7-8 per group, average of 72 hours, Two-way ANOVA). B) The energy expenditure (EE) was examined for Adipoq-*Acbp^-/-^* and control mice housed individually at room temperature (22°C) for 72 hours in metabolic cages. Data are presented as mean of individuals in each group ± SEM (n=7-8 per group, average of 72 hours, Two-way ANOVA). C) The food intake was examined for Adipoq-*Acbp^-/-^* and control mice housed individually at room temperature (22°C) for 72 hours in metabolic cages. Data are presented as mean of individuals in each group ± SEM (n=7-8 per group, average of 72 hours, Two-way ANOVA). D) The water intake was examined for Adipoq-*Acbp^-/-^* and control mice housed individually at room temperature (22°C) for 72 hours in metabolic cages. Data are presented as mean of individuals in each group ± SEM (n=7-8 per group, average of 72 hours, Two-way ANOVA) E) The respiratory exchange ratio (RER) was examined for Adipoq-*Acbp^-/-^*and control mice housed individually at cold (4°C) for 72 hours in metabolic cages. Data are presented as mean of individuals in each group ± SEM (n=7-8 per group, average of 72 hours, Two-way ANOVA). F) The energy expenditure (EE) was examined for Adipoq-*Acbp^-/-^* and control mice housed individually at cold (4°C) for 72 hours in metabolic cages. Data are presented as mean of individuals in each group ± SEM (n=7-8 per group, average of 72 hours, Two-way ANOVA). G) The food intake was examined for Adipoq-*Acbp^-/-^* and control mice housed individually at cold (4°C) for 72 hours in metabolic cages. Data are presented as mean of individuals in each group ± SEM (n=7-8 per group, average of 72 hours, Two-way ANOVA). H) The water intake was examined for Adipoq-*Acbp^-/-^* and control mice housed individually at cold (4°C) for 72 hours in metabolic cages. Data are presented as mean of individuals in each group ± SEM (n=7-8 per group, average of 72 hours, Two-way ANOVA). I) The respiratory exchange ratio (RER) was examined for Adipoq-*Acbp^-/-^*and control mice housed individually at thermoneutrality (30°C) for 72 hours in metabolic cages. Data are presented as mean of individuals in each group ± SEM (n=7-8 per group, average of 72 hours, Two-way ANOVA). J) The energy expenditure (EE) was examined for Adipoq-*Acbp^-/-^* and control mice housed individually at thermoneutrality (30°C) for 72 hours in metabolic cages. All mice were sacrificed by cervical dislocation. Data are presented as mean of individuals in each group ± SEM (n=7-8 per group, average of 72 hours, Two-way ANOVA). K) The food intake was examined for Adipoq-*Acbp^-/-^*and control mice housed individually at thermoneutrality (30°C) for 72 hours in metabolic cages. Data are presented as mean of individuals in each group ± SEM (n=7-8 per group, average of 72 hours, Two-way ANOVA). L) The water intake was examined for Adipoq-*Acbp^-/-^*and control mice housed individually at thermoneutrality (30°C) for 72 hours in metabolic cages. Data are presented as mean of individuals in each group ± SEM (n=7-8 per group, average of 72 hours, Two-way ANOVA). Mice were 9-14 weeks old. All mice were sacrificed by cervical dislocation.

**Figure 6.**
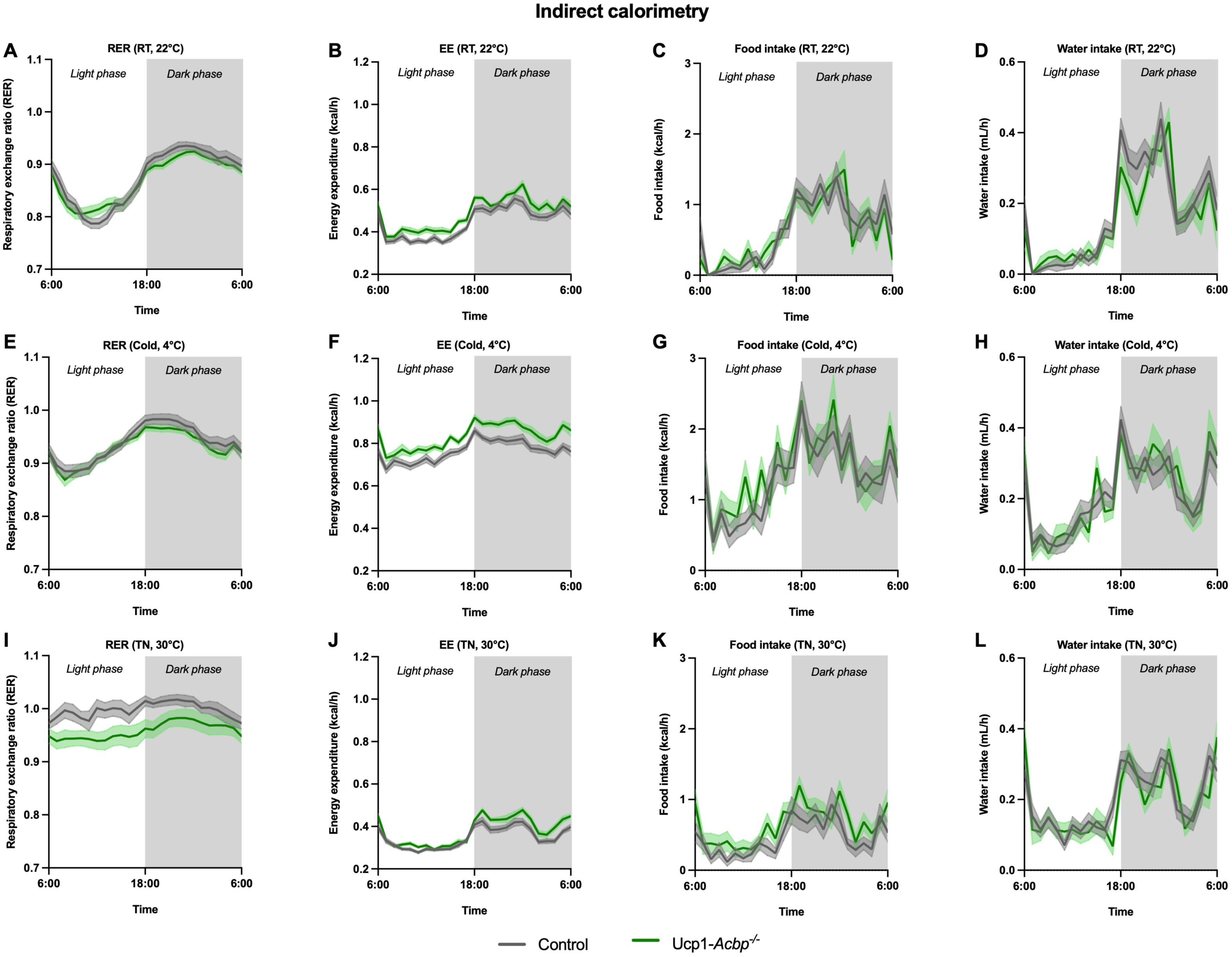
ACBP in adipose tissue is dispensable for systemic energy metabolism in Ucp1-*Acbp^-/-^* mice. A) The respiratory exchange ratio (RER) was examined for Ucp1-*Acbp^-/-^* and control mice housed individually at room temperature (22°C) for 72 hours in metabolic cages. Data are presented as mean of individuals in each group ± SEM (n=7-8 per group, average of 72 hours, Two-way ANOVA). B) The energy expenditure (EE) was examined for Ucp1-*Acbp^-/-^* and control mice (weeks) housed individually at room temperature (22°C) for 72 hours in metabolic cages. Data are presented as mean of individuals in each group ± SEM (n=7-8 per group, average of 72 hours, Two-way ANOVA). C) The food intake was examined for Ucp1-*Acbp^-/-^*and control mice housed individually at room temperature (22°C) for 72 hours in metabolic cages. Data are presented as mean of individuals in each group ± SEM (n=7-8 per group, average of 72 hours, Two-way ANOVA). D) The water intake was examined for Ucp1-*Acbp^-/-^*and control mice housed individually at room temperature (22°C) for 72 hours in metabolic cages. Data are presented as mean of individuals in each group ± SEM (n=7-8 per group, average of 72 hours, Two-way ANOVA). E) The respiratory exchange ratio (RER) was examined for Ucp1-*Acbp^-/-^* and control mice housed individually at cold (4°C) for 72 hours in metabolic cages. Data are presented as mean of individuals in each group ± SEM (n=7-8 per group, average of 72 hours, Two-way ANOVA). F) The energy expenditure (EE) was examined for Ucp1-*Acbp^-/-^* and control mice housed individually at cold (4°C) for 72 hours in metabolic cages. Data are presented as mean of individuals in each group ± SEM (n=7-8 per group, average of 72 hours, Two-way ANOVA). G) The food intake was examined for Ucp1-*Acbp^-/-^* and control mice housed individually at cold (4°C) for 72 hours in metabolic cages. Data are presented as mean of individuals in each group ± SEM (n=7-8 per group, average of 72 hours, Two-way ANOVA). H) The water intake was examined for Ucp1-*Acbp^-/-^*and control mice housed individually at cold (4°C) for 72 hours in metabolic cages. Data are presented as mean of individuals in each group ± SEM (n=7-8 per group, average of 72 hours, Two-way ANOVA). I) The respiratory exchange ratio (RER) was examined for Ucp1-*Acbp^-/-^* and control mice housed individually at thermoneutrality (30°C) for 72 hours in metabolic cages. Data are presented as mean of individuals in each group ± SEM (n=7-8 per group, average of 72 hours, Two-way ANOVA). J) The energy expenditure (EE) was examined for Ucp1-*Acbp^-/-^* and control mice housed individually at thermoneutrality (30°C) for 72 hours in metabolic cages. Data are presented as mean of individuals in each group ± SEM (n=7-8 per group, average of 72 hours, Two-way ANOVA). K) The food intake was examined for Ucp1-*Acbp^-/-^*and control mice housed individually at thermoneutrality (30°C) for 72 hours in metabolic cages. Data are presented as mean of individuals in each group ± SEM (n=7-8 per group, average of 72 hours, Two-way ANOVA). L) The water intake was examined for Ucp1-*Acbp^-/-^*and control mice housed individually at thermoneutrality (30°C) for 72 hours in metabolic cages. Data are presented as mean of individuals in each group ± SEM (n=7-8 per group, average of 72 hours, Two-way ANOVA). Mice were 9-11 weeks old. All mice were sacrificed by cervical dislocation.

### Loss of ACBP in adipose tissue alter acyl-carnitine metabolism in BAT

Despite that loss of ACBP in adipose tissues does not affect systemic energy expenditure, we envisioned that ACBP could play a role in intracellular lipid metabolism in adipocytes. We therefore examined the lipid composition of adipose tissues in both tissue-specific ACBP deficient mouse models using liquid chromatography – mass spectrometry (LC-MS) based lipidomics. Interestingly, we found several acyl-carnitine species to be elevated in BAT from the Adipoq-*Acbp^-/-^* mice housed at RT compared with their control littermates (Figure 7A). Yet, we observed no significant changes in acyl-carnitine levels in BAT from the Adipoq-*Acbp^-/-^* mice housed at 4°C for 7 days (Figure 7B) or in BAT from Ucp1-*Acbp^-/-^*mice housed at RT or 4°C for 7 days (Figure 7C-D). However, lipidomic analyses demonstrated that removal of ACBP from the adipose tissue did not induce any major changes in the lipidome of iWAT from Adipoq-*Acbp^-/-^* mice as well as Ucp1-*Acbp^-/-^* mice independent of housing temperature (Figure S3). Taken together, these data revealed that removal of ACBP from the adipose tissue only induce subtle effects in the lipidomes of WAT and BAT, primarily affecting the composition of the acyl-carnitine pool in BAT from Adipoq-*Acbp^-/-^*mice when housed room temperature.

**Figure 7.**
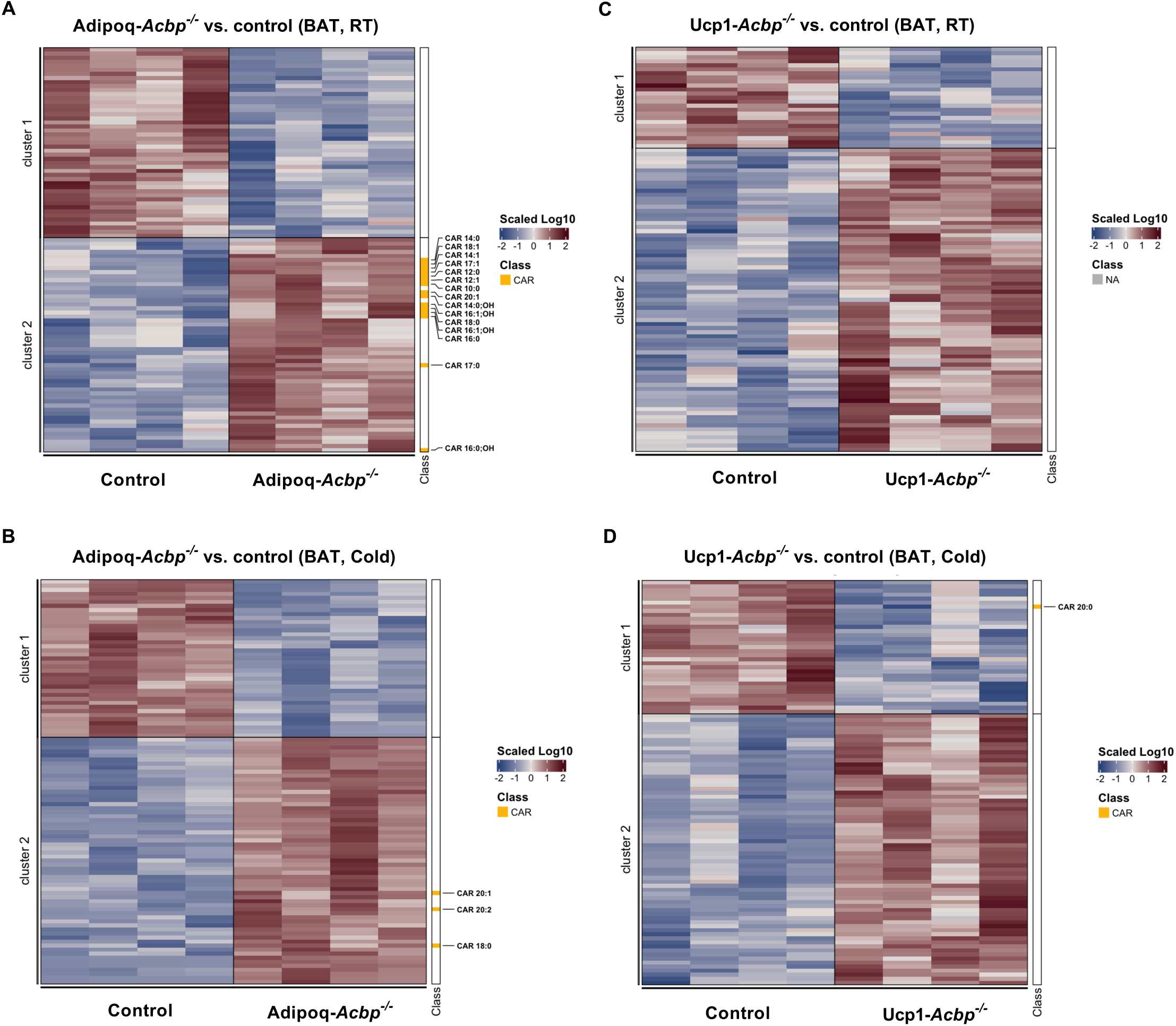
Loss of ACBP in adipose tissue alter acyl-carnitine metabolism in BAT. A) Heatmap of the 100 most regulated lipid metabolites in BAT harvested from Adipoq-*Acbp^-/-^* mice housed individually at room temperature (22°C) for 7 days compared to BAT harvested from control mice housed individually at room temperature (22°C) for 7 days (n=4 per group). B) Heatmap of the 100 most regulated lipid metabolites in BAT harvested from Adipoq-*Acbp^-/-^* mice housed individually at cold (4°C) for 7 days compared to BAT harvested from 9-10 weeks old control mice housed individually at cold (4°C) for 7 days (n=4 per group). C) Heatmap of the 100 most regulated lipid metabolites in BAT harvested from Ucp1-*Acbp^-/-^* mice housed individually at cold room temperature (22°C) for 7 days compared to BAT harvested from 10-11 weeks old control mice housed individually at room temperature (22°C) for 7 days (n=4 per group). D) Heatmap of the 100 most regulated lipid metabolites in BAT harvested from Ucp1-*Acbp^-/-^* mice housed individually at cold (4°C) for 7 days compared to BAT harvested from 11-12 weeks old control mice housed individually at cold (4°C) for 7 days (n=4 per group). Mice were 9-12 weeks old. All mice were sacrificed by cervical dislocation.

This led us to speculate whether increased levels of acyl-carnitines in BAT might indicate that increased levels of fatty acids could be channeled towards mitochondrial fatty acid oxidation e.g. due to increased lipolytic activity in WAT. Thus, we determined the levels of glycerol and non-esterified fatty acids (NEFAs) released from iWAT isolated from control and Adipoq-*Acbp^-/-^* mice housed at RT. We found no significant changes in neither glycerol (Figure 8A) nor NEFA levels (Figure 8B) released from iWAT between the control and Adipoq-*Acbp^-/-^* mice, indicating that ACBP ablation from adipose tissues does not affect the lipolytic activity. Additionally, we found no differences in the levels of glycerol, NEFA and triacylglycerols (TAGs) in blood plasma collected from Adipoq-*Acbp^-/-^* and Ucp1-*Acbp^-/-^* mice housed at either RT or 4°C for 7 days when compared with their respective control littermates (Figure 8C-H).

**Figure 8.**
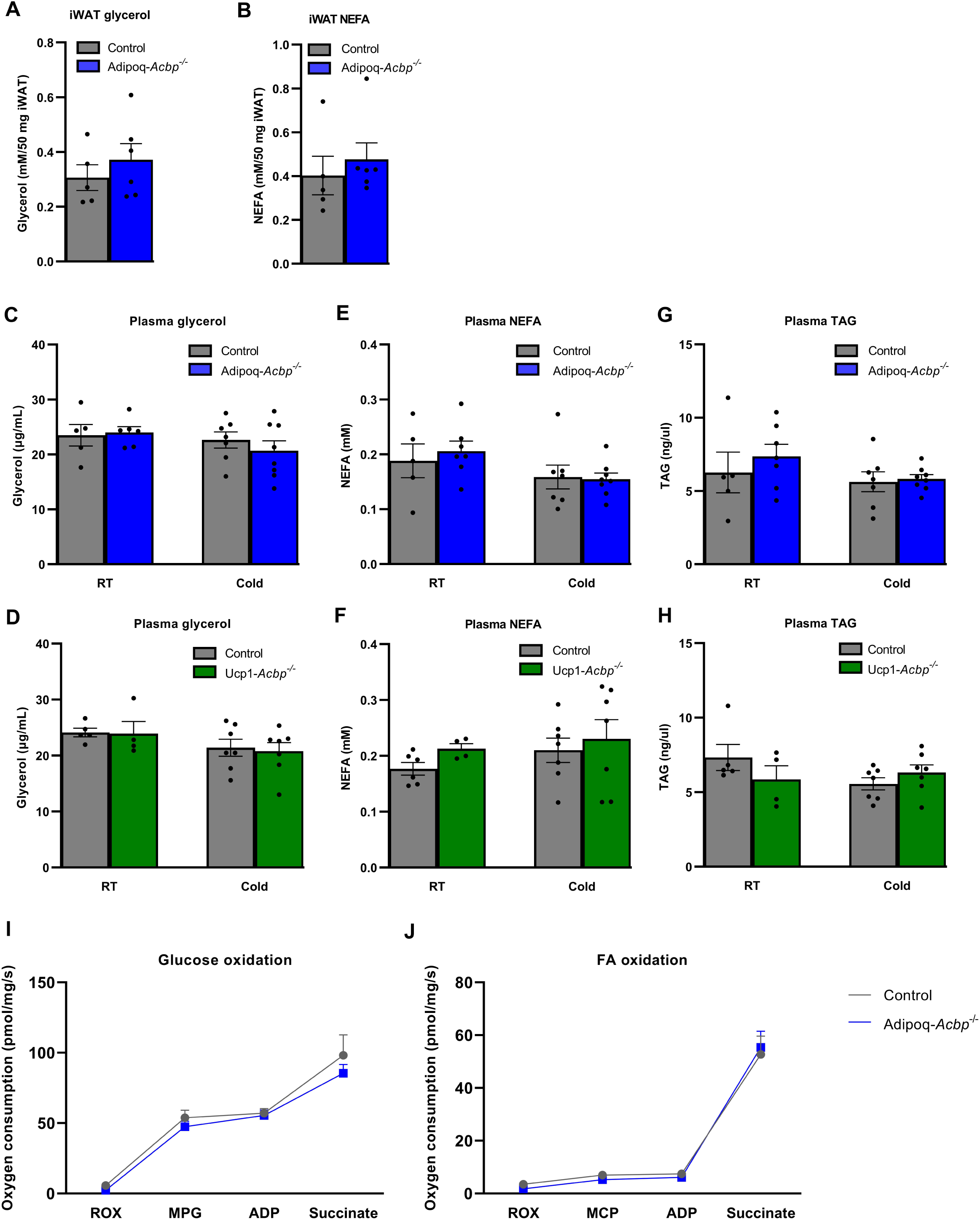
Substrate metabolite levels and mitochondrial respiration is unaffected by ablation of ACBP in adipose tissue. A) Glycerol levels released from iWAT isolated from 13 weeks old Adipoq-*Acbp^-/-^* and control mice were measured by performing *ex vivo* lipolysis. All mice were housed individually at 22°C. Data are presented as mean ± SEM (n=5-6 per group, unpaired t-test). B) Non-esterified fatty acid (NEFA) levels released from iWAT isolated from 13 weeks old Adipoq-*Acbp^-/-^*and control mice were measured by performing *ex vivo* lipolysis. All mice were housed individually at 22°C and sacrificed by cervical dislocation. Data are presented as mean ± SEM (n=5-6 per group, unpaired t-test). C) Plasma glycerol levels were measured for Adipoq-*Acbp^-/-^* and control mice (8-13 weeks old) housed individually at either 22°C or 4°C for 7 days. Data are presented as mean ± SEM (n=5-8 per group, Two-way ANOVA). D) Plasma glycerol levels were measured for Ucp1-*Acbp^-/-^* and control mice (9-11 weeks old) housed individually at either 22°C or 4°C for 7 days. Data are presented as mean ± SEM (n=5-8 per group, Two-way ANOVA). E) Plasma non-esterified fatty acid (NEFA) levels were measured for Adipoq-*Acbp^-/-^* and control mice (8-13 weeks old) housed individually at either 22°C or 4°C for 7 days. Data are presented as mean ± SEM (n=5-8 per group, Two-way ANOVA). F) Plasma non-esterified fatty acid (NEFA) levels were measured for Ucp1-*Acbp^-/-^*and control mice (9-11 weeks old) housed individually at either 22°C or 4°C for 7 days. Data are presented as mean ± SEM (n=5-8 per group, Two-way ANOVA). G) Plasma triacylglycerol (TAG) levels were measured for Adipoq-*Acbp^-/-^* and control mice (8-13 weeks old) housed individually at either 22°C or 4°C for 7 days. Data are presented as mean ± SEM (n=5-8 per group, Two-way ANOVA). H) Plasma triacylglycerol (TAG) levels were measured for Ucp1-*Acbp^-/-^* and control mice (9-11 weeks old) housed individually at either 22°C or 4°C for 7 days. Data are presented as mean ± SEM (n=5-8 per group, Two-way ANOVA). I) Glucose oxidation for BAT isolated from 12-14 weeks old Adipoq-*Acbp^-/-^* and control mice housed individually at 22°C was measured using Oroboros Oxygraph-2K instruments. Data are presented as mean ± SEM (n=7-8 per group, Two-way ANOVA). ROX = residual oxygen consumption. MPG = Malate, pyruvate and glutamate. ADP = Adenosine diphosphate. J) Fatty acid oxidation for BAT isolated from 12-14 weeks old Adipoq-*Acbp^-/-^* and control mice housed individually at 22°C was measured using Oroboros Oxygraph-2K instruments. Data are presented as mean ± SEM (n=7-8 per group, Two-way ANOVA). ROX = residual oxygen consumption. MCP = Malate, Co-enzyme A, palmitic acid. ADP = Adenosine diphosphate. In Figure A, B, I and J, mice were sacrificed by cervical dislocation, while in Figure C-H, mice were anesthetized for 5 minutes and subsequently sacrificed by heart puncture to draw blood.

### Mitochondrial respiration is unaffected by ablation of ACBP in adipose tissue

Since removal of ACBP from adipose tissues does not affect the lipolytic activity in iWAT as well as the plasma levels of glycerol, NEFA and TAG, we speculated whether the increased levels of acyl-carnitines in BAT could indicate that ACBP is required to support transport and oxidation of fatty acids in BAT. Thus, we determined oxygen consumption in BAT isolated from Adipoq-*Acbp^-/-^*mice housed at RT using high-resolution respirometry to examine the mitochondrial respiratory capacity. Since we only found differences in acyl-carnitine levels in BAT from the Adipoq-*Acbp^-/-^* mice and not the Ucp1-*Acbp^-/-^* mice, we only examined oxygen consumption in BAT from Adipoq-*Acbp^-/-^*mice. At baseline with no substrates added (ROX), we found no significant differences in oxygen consumption for glucose oxidation in BAT from Adipoq-*Acbp^-/-^* mice (Figure 8I-J). Furthermore, we observed no differences in leak respiration (PMG/MCP), coupled respiration combining oxidative phosphorylation and ATP synthesis (ADP) or maximal coupled respiration (succinate) during either glucose or palmitate oxidation in BAT from Adipoq-*Acbp^-/-^* mice (Figure 8I-J). In summary, our data show that ACBP ablation in adipose tissues does not affect glucose oxidation (Figure 8I) or FA oxidation (Figure 8J) in BAT under basal and non-stimulated conditions.

Collectively, our present study shows that ACBP is dispensable for adipose tissue function in mice.

## Discussion

ACBP is recognized as a key regulator of intracellular lipid metabolism, particularly in facilitating the utilization of long-chain fatty acyl-CoAs in various anabolic and catabolic pathways(10). We have previously shown that systemic loss of *Acbp* in mice as well as specific deletion in keratinocytes results in increased energy expenditure, increased food intake, increased browning of iWAT and resistance to the diabetogenic effects of a high-fat diet(12). Consistently, Joseph et al. reported that tamoxifen-inducible systemic loss of ACBP also prevents high-fat diet induced weight gain in mice(19). Systemic loss of *Acbp* in mice results in a higher respiratory quotient than observed for wild type and keratinocyte-specific *Acbp*^-/-^ mice. This suggests that ACBP is likely to facilitate oxidation of fatty acids in other cell types than keratinocytes and thereby contribute to systemic utilization of fatty acids as energy substrates(12). In the present study, we have established Adipoq-*Acbp^-/-^* mice and Ucp1-*Acbp^-/-^* mice to examine the function of ACBP in BAT and WAT. Despite elaborate analyses, we find that loss of ACBP in adipose tissues have no effect on body- and tissue weights, food intake, systemic energy homeostasis or the ability to maintain body temperature, even after cold exposure.

Intracellular trafficking of acyl-CoA esters between metabolic compartments is essential for maintaining lipid homeostasis and cellular energy production. ACBP facilitates intracellular transport of acyl-CoA esters to key metabolic enzymes, including carnitine palmitoyltransferase 1 (CPT1), the rate-limiting enzyme of mitochondrial fatty acid oxidation (FAO). CPT1, located on the outer mitochondrial membrane, catalyzes the transfer of long-chain acyl groups from CoA to carnitine, enabling their translocation across the inner mitochondrial membrane for subsequent β-oxidation. ACBP enhances the delivery of acyl-CoA esters to CPT1 via direct interaction(20, 21), thereby promoting FAO under conditions of increased energy demand. This function has recently been found particularly important for driving cell survival and proliferation of glioblastoma and non-small cell lung cancer cells(22–25). A similar function could be equally important in adipose tissues with high lipid turnover, where efficient acyl-CoA utilization is crucial for metabolic adaptation. In this study, lipidomic analyses revealed elevated levels of long-chain acyl-carnitines in BAT from Adipoq-*Acbp^-/-^* mice when housed at room temperature (Figure 7A-B). However, we did not observe a similar change in BAT from Ucp1-*Acbp^-/-^* mice (Figure 7C-D). This indicates that there might be an increased flow of fatty acids from WAT to BAT or reduced oxidation of fatty acids in BAT in Adipoq-*Acbp^-/-^* mice. Reduced utilization of fatty acids in BAT is likely to be the cause, as we did not detect any changes in plasma NEFA levels or lipolytic activity in WAT in Adipoq-*Acbp^-/-^* mice relative to control mice (Figure 8A-B and E). Notably, upon cold exposure of Adipoq-*Acbp^-/-^*mice, the difference in acyl-carnitine levels disappears, likely due to a switch towards increased glucose oxidation in both control and Adipoq-*Acbp^-/-^* mice. A similar switch in substrate utilization in BAT upon cold exposure is in line with recent observations(26), yet not sufficient to significantly change the RER of Adipoq-*Acbp^-/-^*mice during cold exposure (Figure 5E).

Intracellular and extracellular sources of fatty acids as well as glucose are commonly known as preferred energy sources to fuel non-shivering thermogenesis in BAT(27–29). To further understand the mechanisms explaining the elevated acyl-carnitines levels in BAT in Adipoq-*Acbp^-/-^* mice, we examined to what extend loss of ACBP affected fatty acid-dependent respiration in BAT using high-resolution respirometry. Despite oxygen consumption in BAT using palmitate as substrate is relatively low, loss of ACBP did not affect respiration in BAT using palmitate as substrate (Figure 8J). Similarly, loss of ACBP did not affect respiration when O_2_ consumption was driven by glycolytic and TCA intermediates (Figure 8I). This observation supports the notion that the mitochondrial respiratory capacity in BAT is intact in the absence of ACBP, despite elevated levels of acyl-carnitines.

Both systemic and keratinocyte-specific ablation of *Acbp* result in increased energy expenditure, increased food intake and increased browning of iWAT, phenotypes that are reversed by blunting β-adrenergic signaling by isopropanol injections or by housing mice at thermoneutrality(12). This and the fact that artificial skin barriers rescue the impaired SREBP-regulated gene expression in livers of *Acbp*^-/-^ mice(30, 31), argue that loss of ACBP in keratinocytes results in an impaired epidermal barrier, which causes systemic metabolic effects. The impaired epidermal barrier is likely to be due to diminished synthesis of ultra very-long and very-long chain ceramides (our unpublished results) as these are essential for epidermal barrier function(32), and as ACBP interacts and potently activates ceramide synthase 2 and 3 (CerS2 and CerS3)(33, 34). While CerS3 is primarily expressed particularly in keratinocytes of the epidermis and spermatogenic cells of the testes, CerS2 is more widely expressed also in white- and brown adipose tissue(35). Therefore, we reasoned that lack of ACBP in the adipose tissue could compromise the composition of ceramide- and sphingolipids of WAT and BAT. However, both our transcriptomics data (Figure 4) and lipidomics data (Figure 7) show no indications of significant changes in expression of ceramide synthases or ceramide composition in BAT and iWAT from either Adipoq-*Acbp^-/-^* mice or Ucp1-*Acbp^-/-^* mice. This indicates that ACBP is not likely to play a vital role for ceramide synthase activity and ceramide synthesis in adipose tissues.

The lack of profound metabolic phenotypes in both Adipoq-*Acbp^-/-^*and Ucp1-*Acbp^-/-^* mice raises intriguing questions about the presence of compensatory mechanisms within adipose tissues that may mitigate the loss of ACBP, such as metabolic rewiring by other acyl-CoA binding proteins e.g. acyl-CoA binding domain proteins, sterol-carrier protein 2 or fatty acid binding protein 1(10, 36). However, our transcriptomic data (Figure 4) does not show transcripts encoding such proteins being upregulated in the absence of ACBP. The results presented in this study challenge our prior assumptions regarding an indispensable role of ACBP in adipose tissue function and metabolism. Given the noticeable effects on systemic energy metabolism observed in full-body ACBP KO mice(12) in addition to the known role of ACBP in facilitating fatty acid oxidation, future studies will uncover whether more oxidative tissues, like skeletal muscle and heart, exhibit a stronger dependency on ACBP for metabolic homeostasis.

## Materials and Methods

### Experimental model

Male mice with conditional targeting of the *Acbp* gene (*Acbp^flox/flox^*) in adipose tissue (Adipoq-*Acbp^-/-^*) or brown adipose tissue (Ucp1-*Acbp^-/-^*) were generated as previously described(17, 37). Ucp1-Cre and Adipoq-Cre mice were obtained from The Jackson Laboratories (B6.FVB-Tg(Ucp1-Cre)1Evdr/J strain 024670 and B6.FVB-Tg(Adipoq-Cre)1Evdr/J strain 028020). *Acbp^flox/flox^*littermates not carrying the Cre allele were used as controls. Unless otherwise specified, mice were group housed in individually ventilated cages (IVC) (Scanbur A/S, Sealsafe Plus GM500) and subjected to standard housing conditions including a 12-hour light/dark cycle, 55% ± 5% relative humidity and housing temperatures at 22°C ± 3°C, while given ad libitum access to chow food (Altromin 1324) and water. All methods were carried out in accordance with relevant guidelines and regulations and were reported in accordance with the ARRIVE guidelines. All experiments involving animals were approved by the Danish Animal Experiment Inspectorate.

### Room temperature and cold exposure experiments

Prior to all experiments, male mice were housed individually in IVC cages for 7 days at standard housing conditions for acclimatization, unless otherwise noted. Hereafter, the mice were housed at either room temperature (RT, 22°C) or 4°C (cold) for 7 days with ad libitum access to food and water. Body weight and food intake were monitored. Body composition was examined using TD-NMR (MiniSpec LF50 whole body composition analyzer). Body temperature was monitored by implantation of subcutaneous transponders (Avidity Science, BMDS IPTT-300). To monitor body temperature mice (10-13 weeks) were housed at RT from hour 0 to 24 and hereafter at 4°C from hour 24 and until the experiment was terminated. The body temperature was recorded at hour 0, 24, 28, 36, 48 and 72 after starting the experiment. Food was removed during the last 6 hours of the experiment. All mice were fasted two hours prior to sacrifice. Depending on the following experiments mice were sacrificed by either cervical dislocation or anesthesia for 5 minutes followed by heart puncture to draw blood. Anesthesia used: 25% hypnorm (0.315 mg fentanylcitrate, 10 mg of fluanisone, 0.5 mg methylparahydroxybenzoate and 1 mL sterile milliQ water) and 25% midazolam (Midazolam, 5 mg/mL) in sterile milliQ water, dosage: 0.01 mL/g mouse. EDTA plasma and tissues of interest were harvested and snap frozen in liquid nitrogen.

### Isolation of stroma vascular fractions and differentiation to mature adipocytes

Stroma vascular fractions were isolated from intrascapular BAT and inguinal WAT from 6 to 8-week-old *Acbp^-/-^* male mice and induced for adipogenic differentiation as described(38).

### Quantitative real-time PCR

Tissues and cells were homogenized in TRI Reagent (Sigma) and total RNA was extracted in accordance with the manufacturer’s protocol. RNA concentration and quality was evaluated using a NanoDrop^TM^ 2000C spectrophotometer (Thermo Scientific) and by agarose gel electrophoresis. Prior to cDNA synthesis 1 μg RNA was treated with 10U/μL DNase I recombinant (Roche) and RNA was reverse transcribed into cDNA as previously described(30). mRNA expression levels were analyzed by real-time quantitative PCR (RT-qPCR) using BIO-RAD CFX Opus 384® and normalized to *Tfiib* expression. Primers used: *Tfiib* forward primer 5′-GTTCTGCTCCAACCTTTGCCT, *Tfiib* reverse primer 5’-TGTGTAGCTGCCATCTGCACTT, *Acbp* forward primer 5’-TTTCGGCATCCGTATCACCT, *Acbp* reverse primer 5’-TTTGTCAAATTCAGCCTGAGACA-3′ *β-actin* forward 5’ - TGT TAC CAA CTG GGA CGA CA, *β-actin* reverse 5’ - GGG GTG TTG AAG GTC TCA AA, Pparγ forward 5’ – CCC TGG CAA AGC ATT TGT AT, Pparγ reverse 5’ - GAA ACT GGC ACC CTT GAA AA, *Fabp4* forward 5’ – CAT CAG CGT AAA TGG GGA TT, *Fabp4* reverse 5’ - TCG ACT TTC CAT CCC ACT TC, *Cebpα* forward 5’ - CAA GAA CAG CAA CGA GTA CCG, *Cebpα* reverse 5’ - GTC ACT GGT CAA CTC CAG CAC, *Ucp*1 forward 5’ – CCT TCC CGC TGG ACA CTG, *Ucp1* 5’ reverse – GGC CTT CAC CTT GGA TCT GA, *Cidea* forward 5’ – AGGGACAGAAATGGACACCG, *Cidea* reverse 5’ – GTTGCTTGCAGACTGGGACA, *Ppargc1a* forward 5’ – AAG GTC CCC AGG CAG TAG AT, *Ppargc1a* reverse 5’ - GCG GTA TTC ATC CCT CTT GA.

### 2.4 Western blot analysis

Protein extraction from tissues was conducted as previously described(30). Protein concentration was determined using Pierce^TM^ bicinchoninic acid (BCA) assay (ThermoFisher Scientific) in accordance with the manufacturer’s protocol. Equal amounts of protein were separated on 15% SDS gels by sodium dodecyl sulfate-polyacrylamide gel electrophoresis (SDS-PAGE). Subsequently, proteins were blotted onto an Immun-Blot® polyvinylidene difluoride (PVDF) membrane (BIO-RAD) by semi-dry blotting at 200 V and 36 mA/membrane for 1 hour. Membranes were blocked in 5% non-fat dry milk powder (Natur Drogeriet) dissolved in 1x TBST (10x TBS (Ultra Pure Tris (Invitrogen), NaCl (Sigma) and milliQ water, pH 7.6), milliQ water and 0.1% Tween20 (Sigma)) at RT, 1 hour or overnight (ON) at 4°C, followed by incubation with primary and secondary antibodies diluted in 5% milk in 1x TBST. Antibodies used: rabbit anti-ratACBP (1:2000)(30), mouse anti-TFIIB (1:1000) (Santa Cruz), goat anti-rabbit IgG horse radish peroxidase (HRP) conjugate (1:7500) (Promega), and goat anti-mouse IgG HRP conjugate (1:7500) (Promega). Membranes were incubated with Immubilon® Forte Western HRP Substrate (Millipore) in darkness for 2 minutes and developed with chemiluminescence using an Amersham ImageQuant 800 (GE Healthcare Bio-Science AB) luminescent image analyzer. Uncropped Western blots are shown in Figure S4 and S5.

### Histology

Inguinal white adipose tissue (iWAT) and brown adipose tissue (BAT) from male control, Ucp1-*Acbp^-/-^* and Adipoq-*Acbp^-/-^* mice (8 to 12 weeks) housed at RT or cold for 7 days were isolated and fixed in 3.7% PFA (10x PBS, 37% PFA (Sigma) and milliQ water) at 4°C for 24 hours. All mice were sacrificed by cervical dislocation. Preparation of tissues for paraffin embedding was conducted by dehydrating the tissue in ethanol and xylene. Paraffin blocks were cut in 3.5 μm sections, mounted on microscopic slides and dried at 60°C for 2 hours and subsequently at 37°C ON. Hereafter, paraffin sections were deparaffinized, rehydrated with ethanol and H_2_O before immunohistochemistry (IHC) staining (Department of pathology, OUH, Denmark). Sections were stained with anti-ACBP (1:500) (Cell signaling), which was detected with a VS200 slide scanner (Olympus) and data was processed using OlyVIA 4.1 (Olympia).

### Measurements of TAG, NEFA and glycerol

Triacylglycerides (TAG), non-esterified fatty acids (NEFA) and glycerol levels were evaluated in EDTA plasma samples collected from male control, Ucp1-*Acbp^-/-^* and Adipoq-*Acbp^-/-^* mice (8 to 13 weeks) housed at RT or cold for 7 days. All mice were anesthetized for 5 minutes and subsequently sacrificed by heart puncture to draw blood. Measurements were performed using the Triglyceride Quantification Kit (Sigma), NEFA-HR(2) and NEFA standard kit (Wako) and Glycerol Assay Kit (Abcam) according to the manufacturer’s instructions.

### *Ex vivo* lipolysis

*Ex vivo* lipolysis was performed to measure the NEFA and glycerol levels released from iWAT samples harvested from male control and Adipoq-*Acbp^-/-^* mice (13 weeks) housed at RT. All mice were sacrificed by cervical dislocation. Approximately 2x 50 mg iWAT was harvested, dissected into smaller pieces and transferred to Eppendorf tubes containing 1 mL pre-warmed (37°C) Krebs-Ringer bicarbonate HEPES buffer (pH = 7.4) (Sigma) and 2% fatty acid free BSA (Sigma). 200 μL Krebs-Ringer bicarbonate HEPES buffer was collected as baseline sample after 0 minutes of incubation and snap frozen in N_2_. The tissue samples were incubated shaking (2000 rpm) in Krebs-Ringer bicarbonate HEPES buffer for 30 minutes at 37°C. Hereafter, 200 μL Krebs-Ringer bicarbonate HEPES buffer was collected and snap frozen in N_2_. Finally, the collected samples of buffer was used to measure the NEFA and glycerol levels using the NEFA-HR(2) and NEFA standard kit (Wako) and the Glycerol Assay Kit (Abcam) according to the manufacturer’s instructions.

### Sample extraction for lipidomics

The lipidome of iWAT and BAT harvested from male control, Ucp1-*Acbp^-/-^* and Adipoq-*Acbp^-/-^* mice (9 to 12 weeks) housed at RT or cold for 7 days was examined using liquid chromatography – mass spectrometry (LC-MS) based lipidomics. All mice were sacrificed by cervical dislocation. Approximately 35 mg iWAT and 20 mg BAT tissue was homogenized for 45 seconds in 1 mL ice-cold extraction solvent (1:2 methanol (Biosolve BV)/chloroform (Merck) including the internal standards lipidomics Splashmix (Avanti), ceramide 18:1/17:0 (100 pmol/μL) (Avanti), ^27^D-myristic acid (0.75 μg/μL) (Sigma) and cardiolipin-d5 (100 pmol/μL) (Avanti) using a FastPrep-24^TM^ Classic Instrument (MP Biomedical) homogenizer. LC-MS grade water (Biosolve BV) was added to each sample followed by shaking at 4°C/1000 rpm for 30 minutes and centrifugation at 4°C/16000xg for 10 minutes. The upper aqueous phase (containing metabolites) and the lower organic phase (containing lipids) were transferred to new tubes. The aqueous phase was re-extracted in 350 μL washing solvent (86/14/1 chloroform (Merck)/methanol (Biosolve BV)/LC-MS grade water (Biosolve BV)), incubated at 4°C, 1000 rpm shaking for 20 minutes followed by centrifugation at 4°C/16000xg for 10 minutes. All samples were stored at -20°C. On the day of analysis, the organic phase was dried under a steam of N_2_ and resuspended in 20 μL chloroform/methanol (2:1) followed by incubation at RT, 800 rpm shaking for 5 minutes before centrifugation at RT/16000xg for 5 minutes. The supernatant was transferred to injection vials. Quality control samples (QC) were generated by pooling 5 μL of each sample.

### Liquid chromatography – mass spectrometry (LC-MS) based lipidomics

Samples (0.5 μL) for lipidomics were injected using Vanquish Horizon UPLC (Thermo Fisher Scientific) equipped with an ACQUITY Premier CSH column (2.1 x 100 mm, 1.7 μM, Waters) operated at 55°C. The analytes were eluted at a flow rate of 400 μL/min with the following gradient of eluent A (Acetonitrile/water (60:40), 10 mM ammonium formate, 0.1% formic acid) and eluent B (Isopropanol/acetonitrile (90:10), 10 mM ammonium formate, 0.1% formic acid). The mobile phase gradient was set as follows: from 0-0.5 minutes; 40% eluent B, from 0.5-0.7 minutes; 40-43% eluent B, from 0.7-0.8 minutes; 43-65% eluent B, from 0.8-2.3 minutes; 65-70% eluent B, from 2.3-6 minutes; 70-99% eluent B, from 6-6.8 minutes; 99% eluent B and from 6.8-7 minutes; 99-40% eluent B. The system was then equilibrated for 3 minutes with the initial conditions. The UPLC flow was coupled to a TimsTOF Flex (Bruker) instrument for mass spectrometric analysis in both positive and negative ion modes. Data files were processed with Metaboscape (v 2024B, Bruker). Data were annotated using both an in-built rule-based annotation approach and the LipidBlast MS2 library (LipidBlast)(39). The resulting feature tables were filtered and merged for positive and negative ion modes using MetaboLink(40). Final data were analyzed using MetaboAnalyst(41) (ver. 6.0). Data are reposited at Mendelay doi:10.17632/fxbshhpwh7.1.

### RNA sequencing

RNA was extracted from BAT harvested from control and Ucp1-*Acbp^-/-^* mice (10 to 11 weeks) housed at RT as well as from BAT and iWAT harvested frm control and Adipoq-*Acbp^-/-^* mice (9 to 10 weeks) housed at RT. All mice were sacrificed by cervical dislocation. 500 ng RNA in a final volume of 25 μL DEPC-treated water was prepared and sample preparation was performed as described in the NEBNext Poly(A) mRNA Magnetic Isolation Module kit and NEBNext Ultra II RNA Library Prep Kit E7770 (New England Biolabs, Ipswich, MA, USA). The amplified libraries were validated by Agilent 2100 Bioanalyzer using a DNA 1000 kit (Agilent Technologies, Inc., Santa Clara, CA, USA) and quantified by qPCR using the KaPa Library Kits (KaPa Biosystems, Wilmington, MA, USA) Hereafter, RNA sequencing was performed using the Novaseq 6000 (Illumina). Raw sequencing reads were mapped to the mouse transcriptome using salmon(42) version 1.7.0 with transcript annotation from Ensembl ver. 101(43). Subsequently, gene counts were obtained by merging transcript counts using tximport(44) and differential gene expression analyzed with DESeq2(45) with shrinkage of fold-changes using apeglm(46). Significantly expressed genes were identified by the Wald test using a negative binomial modelling of gene counts. Volcano plots were generated using the EnhancedVolcano R package(47). Data are reposited at ArrayExpress no. E-MTAB-14881.

### Indirect calorimetry

Indirect calorimetry was conducted using the PhenoMaster NG 2.0 Home Cage System (TSE systems, Bad Homburg, Germany). Prior to the experiment, a complete calibration protocol for the gas analysers was run according to the manufacturer’s recommendations, and the mice were weighed. Male control, Ucp1-*Acbp^-/-^* (9-11 weeks) and Adipoq-*Acbp^-/-^* mice (9 to 14 weeks) were housed individually. Mice were subjected to 50% humidity, a 12-hour light/dark cycle and given access to ad libitum food (Altromin 1324) and water. Prior to measurements the mice were allowed to acclimate in the system for 3 days at RT. Hereafter, energy expenditure, respiratory exchange ratio, food and water intake was assessed over a period of 72 hours at RT (22°C), followed by 72 hours at cold (4°C), and finally 72 hours at thermoneutrality (30°C). Upon termination of the experiment, all mice were sacrificed by cervical dislocation. Temperature changes were performed through a 24-hour long ramp between each 72-hour measurement interval. The sampling rate was 60 seconds and datapoints were averaged per hour during data analysis. Data were analyzed as the average of 72 hours and presented as the mean of individuals in each group ± SEM.

### Mitochondrial respiratory measurements

BAT was isolated from male control and Adipoq-*Acbp^-/-^*mice (12 to 14 weeks) housed individually at RT with access to ad libitum chow food (Altromin 1324) and water. All mice were sacrificed by cervical dislocation. The isolated tissue was placed in ice-cold BIOPS buffer (2.77 mM CaK_2_EGTA, 7.23 mM K_2_EGTA, 5.77 mM Na_2_ATP, 6.56 mM MgCl_2_ · 6 H2O, 20 mM Taurine, 15 mM Na_2_Phosphocreatine, 20 mM Imidazole, 0.5 mM Dithiothreitol, 50 mM MES, pH 7.1) and kept on ice while connective tissue and capillaries were removed. Hereafter, BAT was dissected into smaller pieces and moved to ice-cold MiR05 buffer (110 mM sucrose, 60 mM potassium lactobionate, 0.5 mM EGTA, 3 mM MgCl_2_ · 6 H2O, 20 mM taurine, 10 mM KH_2_PO_4_, 20 mM HEPES, 1g/L BSA, pH 7.1) and washed for 10 minutes shaking on ice. BAT was dried briefly on soaking paper, weighted (∼20 mg) and placed in the respirometry chamber (Oxygraph-2k, Oroboros instruments, Innsbruck, Austria) containing MiR05 at 37°C. Subsequently, 2 µM digitonin was added to permeabilize the plasma membrane of the brown adipocytes. Hereafter, either glucose or fatty acid oxidation was evaluated. To examine glucose oxidation: 2 mM Malate, 10 mM glutamate, and 5 mM pyruvate were added to measure state 2 respiration (CI). State 3 respiration with complex I linked substrates were assessed by adding 5 mM ADP (CIp). 0.01 mM cytochrome C was added to control for outer mitochondrial membrane integrity. Hereafter, 10 mM succinate was added to achieve state 3 maximal coupled respiration (CI+IIp). To examine fatty acid oxidation: 2 mM malate, 40 mM CoA and 5 mM palmitic acid were added to measure state 2 respiration (ETF), followed by addition of 5 mM ADP to achieve state 3 respiration (ETFp). 0.01 mM cytochrome C was added to control for outer mitochondrial membrane integrity. Then, 10 mM succinate was added to achieve maximal coupled respiration (CI+IIp) with parallel electron input from complex I and II. Finally, 10 mM etomoxir was added to inhibit fatty acid oxidation via CPT1. The mitochondrial respiratory capacity was measured using high-resolution respirometry and data was recorded and analyzed in Datlab software (Oroboros Instruments, Innsbruck, Austria). All data were normalized to mg of tissue added to the chamber.

### Quantification and statistical analysis

Unless otherwise stated, statistical analysis was performed with GraphPad Prism version 9.5.1 and all data are presented as mean ± SEM with the significance level set to p<0.05. The value of n is included in the figure legends. An unpaired parametric t-test was applied to analyze body weight data, food intake data as well as *ex vivo* lipolysis glycerol and NEFA data. Two-way ANOVA was applied to assess repeated measures data including *Acbp* gene expression data, body composition data, tissue weight relative to body weight data, body temperature data, indirect calorimetry data, plasma glycerol, NEFA and TAG data as well as glucose and fatty acid oxidation data. In addition, energy expenditure, food and water intake data from the indirect calorimetry experiment was analyzed by ANCOVA with bodyweight as covariate. The ANCOVA analysis was provided by the NIDDK Mouse Metabolic Phenotyping Centers (MMPC, www.mmpc.org) using the Energy Expenditure Analysis page (http://www.mmpc.org/shared/regression.aspx).

## Resource availability

RNAseq data are reposited at ArrayExpress no. E-MTAB-14881. Lipidomics sata are reposited at Mendelay doi:10.17632/fxbshhpwh7.1.

## Lead contact

Requests for further information and resources should be directed to lead contact, Prof. Nils J. Færgeman.

## Materials availability

*Acbp^flox/flox^*mice are available from lead author upon request.

## Data and code availability

Data: All data reported in this publication will be shared by the lead contact upon request. Code: This article does not report any original code.

Additional resources: Any additional information required to reanalyze the data reported in this article is available from the lead contact upon request.

## Acknowledgement

This work was supported by The Novo Nordisk Foundation (NNF20OC0064744). R.P was supported by Independent Research Fund Denmark (2032-00062B), M. F. N. was supported by The Leo Foundation (LF-OC-22-001107). Lipidomics work was performed within the Integra Infrastructure supported by the Novo Nordisk Foundation (NNF20OC0061575).

## Author contributions

Conceptualization, N.J.F. and D.N.; methodology, R.P, M.F.N, D.N., J.H., S.L.; experimental work, R.P, M.F.N., D.N., E.S.J., J.H., P.R., S.L., N.J.F.,; data curation, R.N., T.K.D., J.H.; writing – original draft, R.P, M.F.N.; writing – review & editing, R.P, M.F.N., D.N., N.J.F.; visualization, J.F.K., P.R., T.K.D; supervision, N.J.F., S.M., J.W.K., B.S.A.; funding acquisition, N.J.F.

## References

1. Choe, S. S., Huh, J. Y., Hwang, I. J., Kim, J. I., and Kim, J. B. (2016) Adipose Tissue Remodeling: Its Role in Energy Metabolism and Metabolic Disorders Front Endocrinol (Lausanne) 7, 30

2. Sakers, A., De Siqueira, M. K., Seale, P., and Villanueva, C. J. (2022) Adipose-tissue plasticity in health and disease Cell 185, 419–446

3. Giordano, A., Smorlesi, A., Frontini, A., Barbatelli, G., and Cinti, S. (2014) White, brown and pink adipocytes: the extraordinary plasticity of the adipose organ Eur J Endocrinol 170, R159–171

4. Richard, A. J., White, U., Elks, C. M., and Stephens, J. M. (2000) Adipose Tissue: Physiology to Metabolic Dysfunction In Endotext, Feingold KR, Anawalt B, Blackman MR, Boyce A, Chrousos G, Corpas E, et al., eds. South Dartmouth (MA)

5. Wolfrum, C., and Gerhart-Hines, Z. (2022) Fueling the fire of adipose thermogenesis Science 375, 1229–1231

6. Ameer, F., Scandiuzzi, L., Hasnain, S., Kalbacher, H., and Zaidi, N. (2014) De novo lipogenesis in health and disease Metabolism 63, 895–902

7. Farese, R. V., Jr., and Walther, T. C. (2023) Glycerolipid Synthesis and Lipid Droplet Formation in the Endoplasmic Reticulum Cold Spring Harb Perspect Biol 15,

8. Grabner, G. F., Xie, H., Schweiger, M., and Zechner, R. (2021) Lipolysis: cellular mechanisms for lipid mobilization from fat stores Nat Metab 3, 1445–1465

9. Coleman, R. A. (2019) It takes a village: channeling fatty acid metabolism and triacylglycerol formation via protein interactomes J Lipid Res 60, 490–497

10. Neess, D., Bek, S., Engelsby, H., Gallego, S. F., and Færgeman, N. J. (2015) Long-chain acyl-CoA esters in metabolism and signaling: Role of acyl-CoA binding proteins Prog Lipid Res 59, 1–25

11. Faergeman, N. J., Sigurskjold, B. W., Kragelund, B. B., Andersen, K. V., and Knudsen, J. (1996) Thermodynamics of ligand binding to acyl-coenzyme A binding protein studied by titration calorimetry Biochemistry 35, 14118–14126

12. Neess, D., Kruse, V., Marcher, A. B., Wæde, M. R., Vistisen, J., Møller, P. M., et al. (2021) Epidermal Acyl-CoA-binding protein is indispensable for systemic energy homeostasis Mol Metab 44, 101144

13. Guidotti, A., Forchetti, C. M., Corda, M. G., Konkel, D., Bennett, C. D., and Costa, E. (1983) Isolation, characterization, and purification to homogeneity of an endogenous polypeptide with agonistic action on benzodiazepine receptors Proc Natl Acad Sci U S A 80, 3531–3535

14. Rasmussen, J. T., Rosendal, J., and Knudsen, J. (1993) Interaction of acyl-CoA binding protein (ACBP) on processes for which acyl-CoA is a substrate, product or inhibitor Biochem J 292 (Pt 3), 907–913

15. Alquier, T., Christian-Hinman, C. A., Alfonso, J., and Færgeman, N. J. (2021) From benzodiazepines to fatty acids and beyond: revisiting the role of ACBP/DBI Trends Endocrinol Metab 32, 890–903

16. Bloksgaard, M., Bek, S., Marcher, A. B., Neess, D., Brewer, J., Hannibal-Bach, H. K. et al. (2012) The acyl-CoA binding protein is required for normal epidermal barrier function in mice J Lipid Res 53, 2162–2174

17. Neess, D., Bek, S., Bloksgaard, M., Marcher, A. B., Faergeman, N. J., and Mandrup, S. (2013) Delayed hepatic adaptation to weaning in ACBP-/- mice is caused by disruption of the epidermal barrier Cell Rep 5, 1403–1412

18. Yan, M., Wang, S., Fang, S., E., M., and Yu, B. (2024) Impacts of cold exposure on energy metabolism Frigid Zone Medicine 4, 65–71

19. Joseph, A., Chen, H., Anagnostopoulos, G., Montégut, L., Lafarge, A., Motiño, O., et al. (2021) Effects of acyl-coenzyme A binding protein (ACBP)/diazepam-binding inhibitor (DBI) on body mass index Cell Death & Disease 12, 599

20. Rasmussen, J. T., Faergeman, N. J., Kristiansen, K., and Knudsen, J. (1994) Acyl-CoA-binding protein (ACBP) can mediate intermembrane acyl-CoA transport and donate acyl-CoA for beta-oxidation and glycerolipid synthesis Biochem J 299 (Pt 1), 165–170

21. Abo-Hashema, K. A., Cake, M. H., Lukas, M. A., and Knudsen, J. (2001) The interaction of acyl-CoA with acyl-CoA binding protein and carnitine palmitoyltransferase I Int J Biochem Cell Biol 33, 807–815

22. Harris, F. T., Rahman, S. M., Hassanein, M., Qian, J., Hoeksema, M. D., Chen, H. et al. (2014) Acyl-coenzyme A-binding protein regulates Beta-oxidation required for growth and survival of non-small cell lung cancer Cancer Prev Res (Phila) 7, 748–757

23. Duman, C., Yaqubi, K., Hoffmann, A., Acikgöz, A. A., Korshunov, A., Bendszus, M. et al. (2019) Acyl-CoA-Binding Protein Drives Glioblastoma Tumorigenesis by Sustaining Fatty Acid Oxidation Cell Metab 30, 274–289.e275

24. Duman, C., Di Marco, B., Nevedomskaya, E., Ulug, B., Lesche, R., Christian, S., et al. (2023) Targeting fatty acid oxidation via Acyl-CoA binding protein hinders glioblastoma invasion Cell Death Dis 14, 296

25. Teng, H., Hang, Q., Zheng, C., Yan, Y., Liu, S., Zhao, Y., et al. (2025) In vivo CRISPR activation screen identifies acyl-CoA-binding protein as a driver of bone metastasis Sci Transl Med 17, eado7225

26. Bornstein, M. R., Neinast, M. D., Zeng, X., Chu, Q., Axsom, J., Thorsheim, C. et al. (2023) Comprehensive quantification of metabolic flux during acute cold stress in mice Cell Metab 35, 2077–2092 e2076

27. Wang, Z., Wang, Q. A., Liu, Y., and Jiang, L. (2021) Energy metabolism in brown adipose tissue Febs j 288, 3647–3662

28. Shinde, A. B., Song, A., and Wang, Q. A. (2021) Brown Adipose Tissue Heterogeneity, Energy Metabolism, and Beyond Frontiers in Endocrinology 12,

29. Chitraju, C., Fischer, A. W., Farese, R. V., Jr., and Walther, T. C. (2020) Lipid Droplets in Brown Adipose Tissue Are Dispensable for Cold-Induced Thermogenesis Cell Rep 33, 108348

30. Neess, D., Bloksgaard, M., Bek, S., Marcher, A. B., Elle, I. C., Helledie, T. et al. (2011) Disruption of the acyl-CoA-binding protein gene delays hepatic adaptation to metabolic changes at weaning J Biol Chem 286, 3460–3472

31. Neess, D., Bek, S., Bloksgaard, M., Marcher, A. B., Færgeman, N. J., and Mandrup, S. (2013) Delayed hepatic adaptation to weaning in ACBP-/- mice is caused by disruption of the epidermal barrier Cell Rep 5, 1403–1412

32. Jennemann, R., Rabionet, M., Gorgas, K., Epstein, S., Dalpke, A., Rothermel, U. et al. (2012) Loss of ceramide synthase 3 causes lethal skin barrier disruption Hum Mol Genet 21, 586–608

33. Ferreira, N. S., Engelsby, H., Neess, D., Kelly, S. L., Volpert, G., Merrill, A. H. et al. (2017) Regulation of very-long acyl chain ceramide synthesis by acyl-CoA-binding protein J Biol Chem 292, 7588–7597

34. Jennemann, R., Rabionet, M., Gorgas, K., Epstein, S., Dalpke, A., Rothermel, U. et al. (2012) Loss of ceramide synthase 3 causes lethal skin barrier disruption Hum Mol Genet 21, 586–608

35. Park, W. J., and Park, J. W. (2015) The effect of altered sphingolipid acyl chain length on various disease models Biol Chem 396, 693–705

36. Faergeman, N. J., and Knudsen, J. (1997) Role of long-chain fatty acyl-CoA esters in the regulation of metabolism and in cell signalling Biochem J 323 (Pt 1), 1–12

37. Neess, D., Bloksgaard, M., Bek, S., Marcher, A. B., Elle, I. C., Helledie, T. et al. (2011) Disruption of the acyl-CoA-binding protein gene delays hepatic adaptation to metabolic changes at weaning J Biol Chem 286, 3460–3472

38. Khani, S., Topel, H., Kardinal, R., Tavanez, A. R., Josephrajan, A., Larsen, B. D. M. et al. (2024) Cold-induced expression of a truncated adenylyl cyclase 3 acts as rheostat to brown fat function Nat Metab 6, 1053–1075

39. Kind, T., Liu, K. H., Lee, D. Y., DeFelice, B., Meissen, J. K., and Fiehn, O. (2013) LipidBlast in silico tandem mass spectrometry database for lipid identification Nat Methods 10, 755–758

40. Mendes, A., Havelund, J. F., Lemvig, J., Schwämmle, V., and Færgeman, N. J. (2024) MetaboLink: a web application for streamlined processing and analysis of large-scale untargeted metabolomics data Bioinformatics 40,

41. Pang, Z., Lu, Y., Zhou, G., Hui, F., Xu, L., Viau, C. et al. (2024) MetaboAnalyst 6.0: towards a unified platform for metabolomics data processing, analysis and interpretation Nucleic Acids Res 52, W398–W406

42. Patro, R., Duggal, G., Love, M. I., Irizarry, R. A., and Kingsford, C. (2017) Salmon provides fast and bias-aware quantification of transcript expression Nature Methods 14, 417–419

43. Yates, A. D., Achuthan, P., Akanni, W., Allen, J., Allen, J., Alvarez-Jarreta, J., et al. (2020) Ensembl 2020 Nucleic Acids Res 48, D682–d688

44. Soneson, C., Love, M. I., and Robinson, M. D. (2015) Differential analyses for RNA-seq: transcript-level estimates improve gene-level inferences F1000Res 4, 1521

45. Love, M. I., Huber, W., and Anders, S. (2014) Moderated estimation of fold change and dispersion for RNA-seq data with DESeq2 Genome Biology 15, 550

46. Zhu, A., Ibrahim, J. G., and Love, M. I. (2018) Heavy-tailed prior distributions for sequence count data: removing the noise and preserving large differences Bioinformatics 35, 2084–2092

47. Blighe, K., S Rana, and M Lewis (2018) EnhancedVolcano: Publication-ready volcano plots with enhanced colouring and labeling

