## Supplemental Figure 1 for "Acyl-CoA Binding Protein in White and Brown Adipose Tissue is Dispensable for Systemic Energy Metabolism"

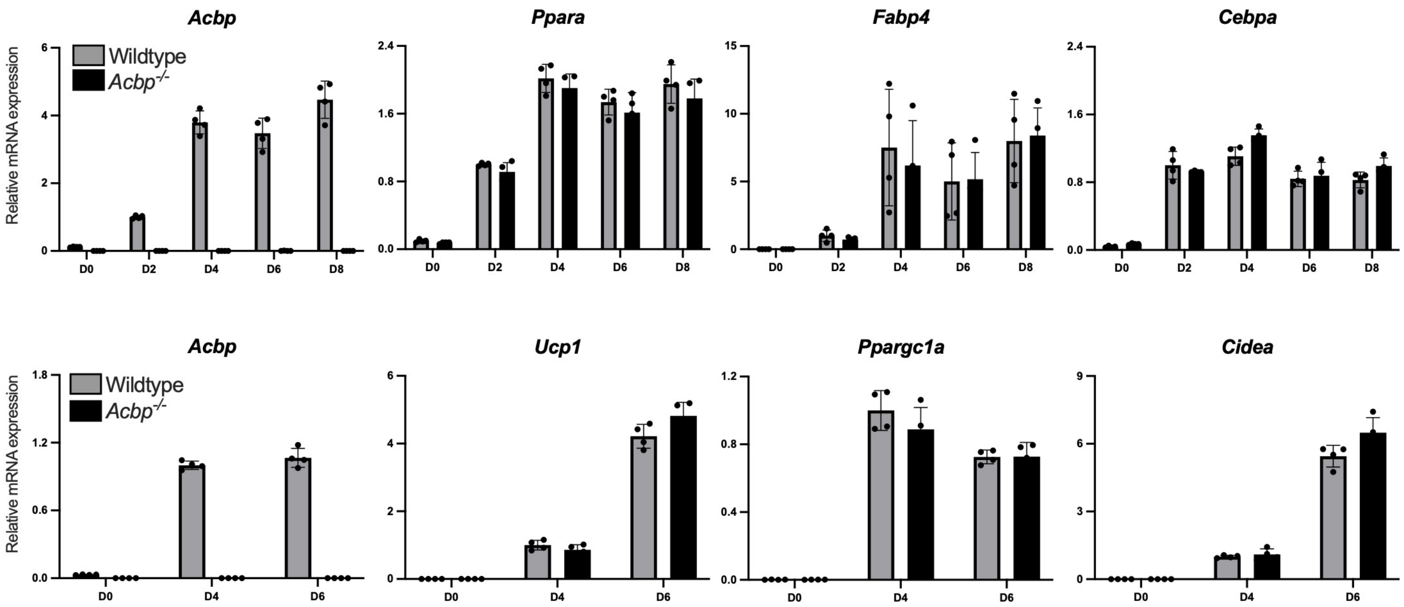


**Figure S1 Relative mRNA expression levels of *Acbp* and adipogenic markers during differentiation of primary preadipocytes isolated from stromal vascular fractions from wildtype and *Acbp*$^{-/-}$ mice.** Stromal vascular fractions (SVF) from of inguinal white adipose tissue (top panel) or brown adipose tissue (bottom panel) was used to isolate primary preadipocytes from wildtype and *Acbp*$^{-/-}$ mice (6-8 weeks). Expression levels of the indicated gene was assesed at D0, D2, D4, D6 and D8 after induction of differentiation and normalized to housekeeping gene *TfIIb*. Relative mRNA expression levels are shown as mean ± SD (n = 4).
