## Supplemental Figure 2 for "Acyl-CoA Binding Protein in White and Brown Adipose Tissue is Dispensable for Systemic Energy Metabolism"

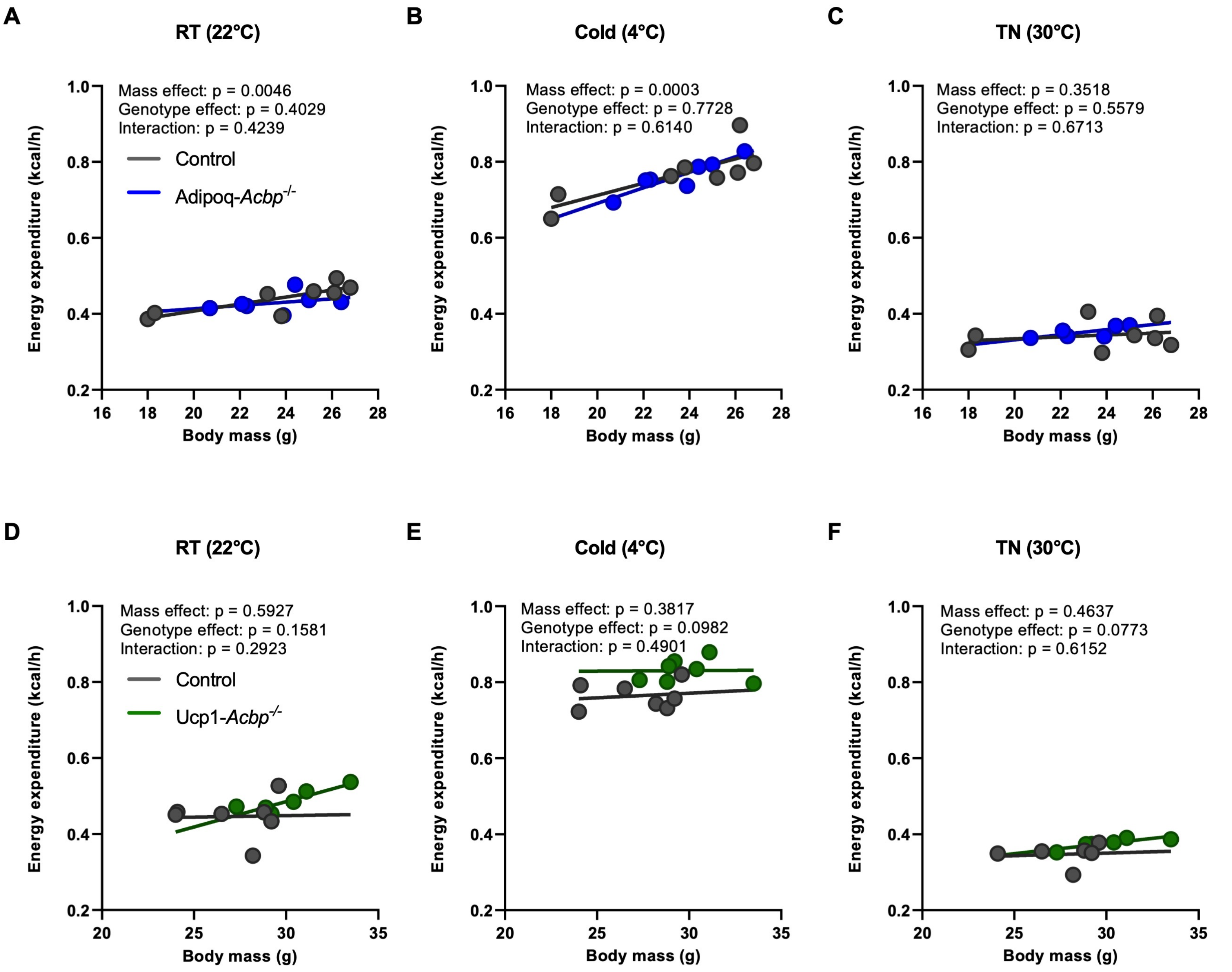


**Figure S2. ACBP in adipose tissue is dispenable for systmeic energy metabolism.** (A-C) Energy expenditure analyzed by ANCOVA with body weight as covariate for 9-14 weeks old control and Adipoq-*Acbp*$^{-/-}$ mice housed individually at A) room temperature (22 °C) for 72 hours, B) cold (4 °C) for 72 hours and C) thermoneutraility (30 °C) for 72 hours. All mice were afterwards sacrificed by cervical dislocation. (n = 7-8 per group). (D-F) Energy expenditure analyzed by ANCOVA with body weight as covariate for 9-11 weeks old control and Ucp1-*Acbp*$^{-/-}$ mice housed individually at D) room temperature (22 °C) for 72 hours, E) cold (4 °C) for 72 hours and F) thermoneutraility (30 °C) for 72 hours. All mice were afterwards sacrificed by cervical dislocation. (n = 7-8 per group).
