## Supplemental Figure 3 for "Acyl-CoA Binding Protein in White and Brown Adipose Tissue is Dispensable for Systemic Energy Metabolism"

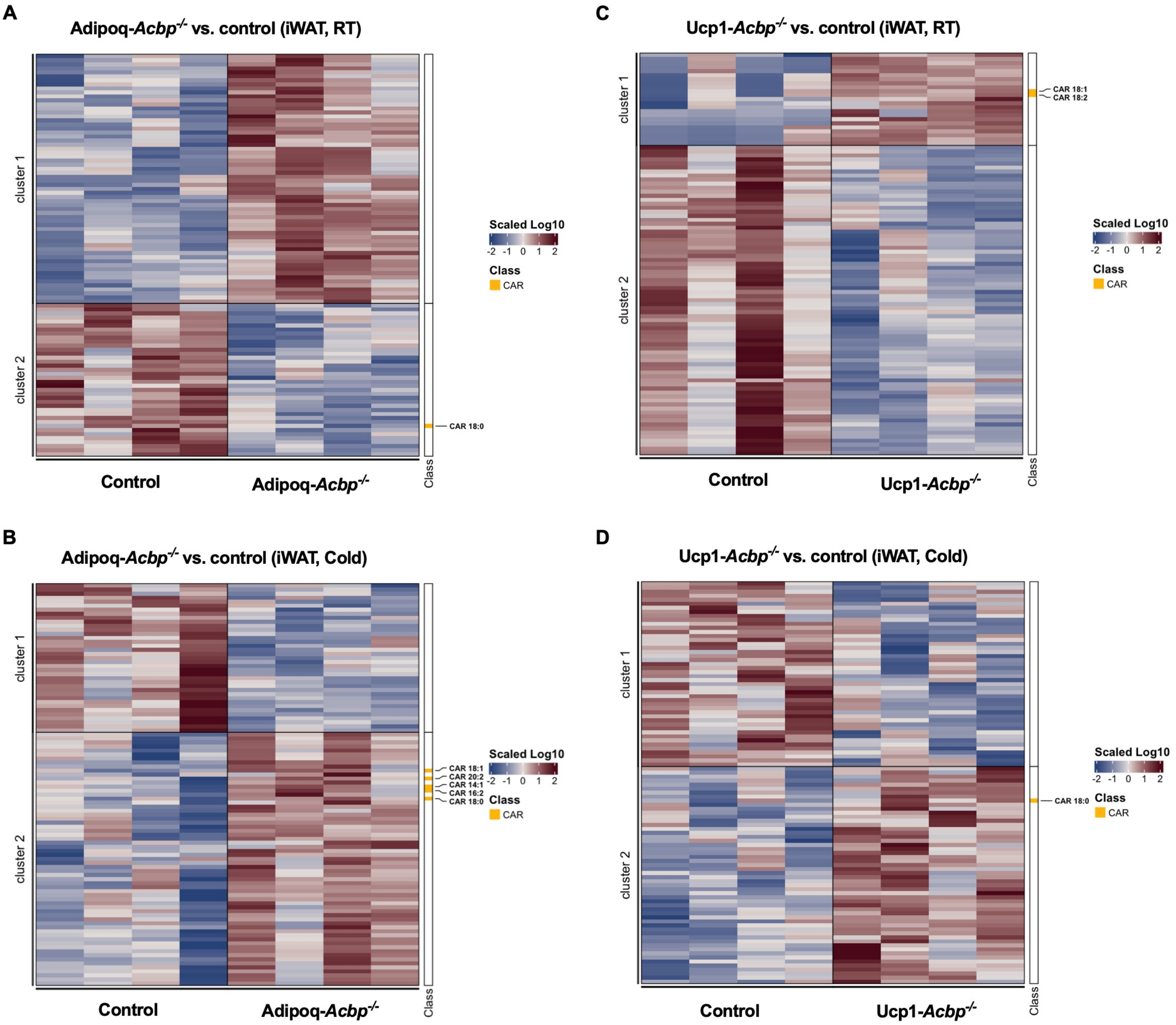


**Figure S3. Loss of ACBP in adipose tissue alter acyl-carnitine metabolism in iWAT.**

A) Heatmap of the 100 most regulated lipid metabolites in iWAT harvested from 9-10 weeks old Adipoq-*Acbp*$^{-/-}$ mice housed individually at room temperature (22°C) for 7 days compared to iWAT harvested from control mice housed individually at room temperature (22°C) for 7 days. All mice were sacrificed by cervical dislocation. (n=4 per group).

B) Heatmap of the 100 most regulated lipid metabolites in iWAT harvested from 9-10 weeks old Adipoq-*Acbp*$^{-/-}$ mice housed individually at cold (4°C) for 7 days compared to iWAT harvested from control mice housed individually at (4°C) for 7 days. All mice were sacrificed by cervical dislocation. (n=4 per group).

C) Heatmap of the 100 most regulated lipid metabolites in iWAT harvested from 10-11 weeks old Ucp1-*Acbp*$^{-/-}$ mice housed individually at room temperature (22°C) for 7 days compared to iWAT harvested from control mice housed individually at room temperature (22°C) for 7 days. All mice were sacrificed by cervical dislocation. (n=4 per group).

D) Heatmap of the 100 most regulated lipid metabolites in iWAT harvested from 10-11 weeks old Ucp1-*Acbp*$^{-/-}$ mice housed individually at cold (4°C) for 7 days compared to iWAT harvested from control mice housed individually at (4°C) for 7 days. All mice were sacrificed by cervical dislocation. (n=4 per group).
